# Single-molecule Imaging of SWI/SNF Chromatin Remodelers Reveal Multi-modal and Cancer-mutant-specific Landscape of DNA-binding Dynamics

**DOI:** 10.1101/2023.07.28.550968

**Authors:** Wilfried Engl, Hendrik Sielaff, Aliz Kunstar-Thomas, Siyi Chen, Woei Shyuan Ng, Ziqing Winston Zhao

## Abstract

Chromatin remodeling, carried out by multi-subunit remodeler complexes, alleviates topological constraints posed by nucleosomes to regulate genome access. Although mutations in the SWI/SNF subfamily of remodelers are implicated in >20% of human cancers, how misregulation of their intranuclear dynamics could underpin cancer remains poorly understood. Combining single-molecule tracking and fluorescence correlation spectroscopy, we probed the live-cell dynamics of three key subunits common to all major human SWI/SNF remodeler complexes (BAF57, BAF155 and BRG1), revealing temporally distinct modes characteristic of free and chromatin-associated diffusion and chromatin-binding. Quantifying residence times of the fully assembled remodeler complex further resolved one transient and two stable binding fractions. Moreover, super-resolved density mapping of single-molecule binding using a newly devised strategy, termed STAR, revealed heterogeneous, nanoscale remodeler binding “hotspots” across the nucleoplasm where multiple binding events preferentially cluster, with particular enrichment of consecutive longer-lived stable binding. Importantly, we showed that the bromodomain plays a key role in modulating the enhancement of remodeler binding dynamics in a DNA-accessibility-dependent manner, but does not facilitate targeting to hyperacetylated chromatin. Finally, we compared the chromatin-binding dynamics of seven common BRG1 mutants implicated in various cancers across tumor types, and uncovered systematic alterations in residence time, binding frequency, fraction of time bound, targeting efficiency and number of binding “hotspots” unique to each point/truncation mutant. Collectively, our findings shed critical insight into the multi-modal landscape regulating the spatio-temporal organizational dynamics of SWI/SNF remodelers to selectively modulate genome accessibility, and could potentially serve as quantitative, mutant-specific signatures for cancers associated with remodeling misregulation.

**SIGNIFICANCE STATEMENT:** Using two complementary approaches, we performed, to our knowledge, the first single-molecule quantification of live-cell dynamics of the fully assembled human SWI/SNF remodeler complex by correlating three key common subunits, and uncovered distinct roles of the bromodomain in modulating chromatin binding/targeting in a DNA-accessibility-dependent manner. Our super-resolved mapping of chromatin-binding also revealed intranuclear “hotspots” where remodelers bind repeatedly in nanometer-scale clusters, as a potential strategy to promote remodeling at these loci. By leveraging previously under-explored parameters, our findings revealed a broader and multi-modal landscape that regulates SWI/SNF-mediated remodeling dynamics in space and time, and established the biophysical basis for aberrant remodeler–chromatin interactions associated with seven mutants implicated in various cancers, which could potentially serve as their unique identifying yardsticks.

## INTRODUCTION

In eukaryotic cells, the genome is packaged inside the nucleus as nucleosomes, each consisting of a core of 147 base pairs of DNA wrapped around a histone octamer, linker histone H1, and a short linker DNA (1). Such structural organization poses a topological challenge when access to the underlying naked DNA is required for various chromatin-based molecular transactions (*e.g.* transcription, DNA repair, replication), and can be alleviated via the process of chromatin remodeling carried out by ATP-dependent, multi-subunit chromatin remodeler complexes (2). Among the different subfamilies of chromatin remodelers, the SWI/SNF (switch/sucrose non-fermentable) remodelers provide DNA access by repositioning or ejecting nucleosomes or evicting histone dimers (3, 4). In mammalian cells, the homolog mSWI/SNF remodeler complexes can be further divided into BAF (canonical BRG1/BRM-associated factors), PBAF (polybromo-associated BAF) and ncBAF (non-canonical BAF) subtypes, each consisting of 8–15 subunits with a total molecular weight (MW) of ∼1–1.5 MDa (5). Structurally, the nucleosome is clamped from both sides in nucleosome-bound SWI/SNF remodeler complexes, with BRG1 being the core ATPase/translocase subunit that targets, binds and anchors the nucleosome with its nucleosome-facing C-terminus, while the N-terminus is anchored in the complex in combination with scaffold subunits such as BAF57 and BAF155 (6–9). Such multi-subunit composition also allows the cell to flexibly assemble remodeler complexes with different tissue- and developmental stage-specific compositions to regulate gene expression and other related processes where and when needed (4, 10). Importantly, mutations in the 29 genes encoding these remodeler subunits (11, 12) have been associated with ∼20% of all human cancers across a wide range of tumor types (13, 14), which is not surprising given that the capability to alter genome organization, accessibility and expression has long been known as a fundamental “enabling characteristic” of cancer (15).

In contrast to our detailed knowledge about the biochemistry, structure and genetics of chromatin remodelers, their spatio-temporal organization and dynamics in live cells are much less understood from a quantitative perspective. Among the few imaging-based studies conducted previously, a series of fluorescence correlation spectroscopy (FCS)- and fluorescence recovery after photobleaching (FRAP)-based studies on the human ISWI remodeler complex have shown that the majority of ISWI remodeler constantly samples nucleosomes via transient binding while a small fraction stably binds to chromatin to carry out remodeling (16, 17). Such a dynamic model was further corroborated and extended to six different remodeler subfamilies (including SWI/SNF) in live yeast cells using single-molecule tracking (SMT) (18). More recently, the dynamic targeting to chromatin of BAF180, a subunit unique to the PBAF subtype of SWI/SNF remodeler complexes, was probed in human cells with SMT and found to occur in a bromodomain-dependent manner (19). However, to date no systematic quantification of the intranuclear dynamics of the human SWI/SNF remodeler subfamily as a whole has been undertaken, nor do we know how misregulation of such dynamics in both space and time might underpin the various types of cancer in which they are implicated.

Herein, we combine SMT, FCS and a newly devised strategy (termed STAR) for super-resolved mapping of intranuclear binding “hotspots” to quantify the live-cell diffusion and chromatin-binding dynamics of three key human SWI/SNF remodeler subunits, namely BAF57, BAF155 and BRG1 (encoded by the SMARCC1, SMARCE1 and SMARCA4 genes, respectively). All three subunits not only perform key catalytic/structural roles (BRG1 is the core ATPase/translocase that directly interacts with the nucleosome, while BAF57 and BAF155 form the backbone of the complex and are critically involved in the complex assembly process), they are also common to all major subtypes of human SWI/SNF remodeler complexes, hence allowing us to reveal dynamic properties that are generic to the entire SWI/SNF remodeler subfamily. These three subunits are also incorporated into the SWI/SNF complex across different stages of the assembly process, with BAF155 being the first, followed by BAF57, and BRG1 being the final subunit to be added (5), thereby allowing us to pinpoint the dynamics that are specific to the fully assembled remodeler complex. Moreover, given that some of the most frequent mutations in human cancers are found in genes encoding the SWI/SNF remodeler subunits (with BRG1 being one of the most frequently mutated) (20, 21), we further probed the chromatin-binding dynamics of seven common mutants of BRG1 implicated in various cancers across tumor types. Our quantitative characterizations across a diverse range of spatio-temporal parameters not only shed key mechanistic insight into the DNA-accessibility-dependent chromatin-binding/targeting by SWI/SNF remodelers, but also uncovered systematic alterations unique to each of the cancer mutants examined, thereby revealing a broad, multi-modal landscape at work for the (mis)regulation of SWI/SNF-mediated remodeling dynamics.

## RESULTS

### Distinct modes of intranuclear dynamics of SWI/SNF remodelers resolved by live-cell FCS

We first performed intranuclear FCS measurements in live HeLa cells expressing each of three remodeler subunits C-terminally fused to mEmerald, while simultaneously monitoring the DNA background stained with Hoechst 33342 (**Fig. 1A**). We found that most of our FCS curves (**Fig. 1B**) were best fitted with a two-component model (*i.e.* two independent diffusing fractions) (**Fig. S1**), with the fast fraction exhibiting a narrow distribution with a mean diffusion coefficient (*D*_fast_) in the range of 13.2–19.7 µm^2^s^-1^ and the slow fraction exhibiting a much broader distribution with a mean diffusion coefficient (*D*_slow_) in the range of 0.29–0.47 µm^2^s^-1^ (**Fig. 1C** and **Table 1**). The fast fraction likely corresponds to the diffusion of the unincorporated individual subunits, as evidenced by the dependence of their diffusion coefficients on their MWs according to a power-law relation derived from the Stokes-Einstein equation, and in line with the diffusion coefficients from previous intranuclear FCS measurements performed in live human cells on a variety of proteins across a wide range of MWs (**Fig. S2**). The slow fraction, however, spans two orders of magnitude and likely corresponds to a mixture of different dynamic behaviors, including diffusion of both the partially and fully assembled remodeler complex as well as chromatin-binding. (For comparison, the diffusion coefficient of an inert 0.58 MDa dextran freely diffusing in the cell nucleus has been found to be ∼1 µm^2^s^-1^ (22), whereas that of the mostly immobile, DNA-bound histone H2B is on the order of ∼0.1 µm^2^s^-1^ (23).)

**Figure 1.**
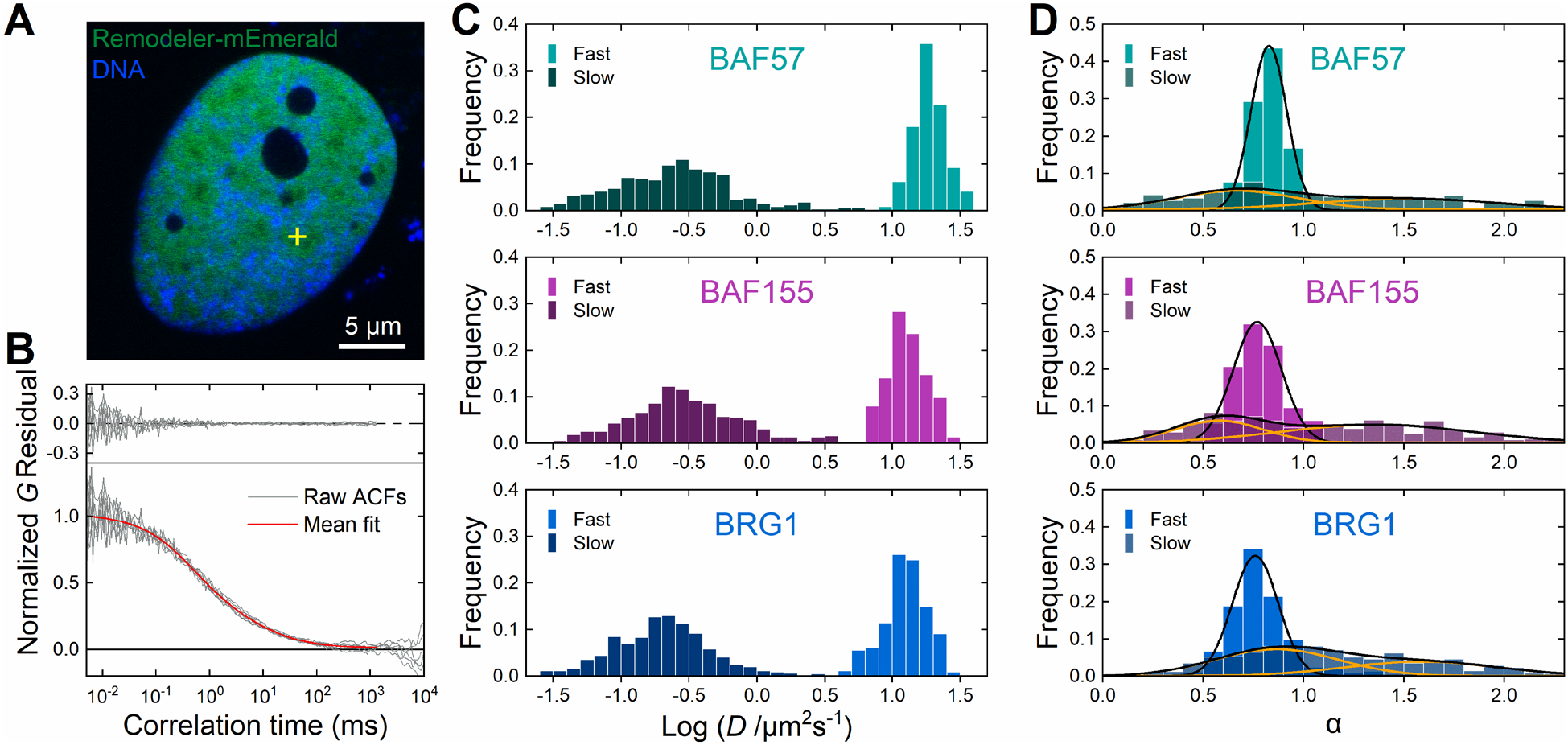
Live-cell FCS resolves distinct modes of intranuclear dynamics of SWI/SNF chromatin remodelers. (**A**) Confocal image of a live HeLa cell nucleus showing a mEmerald-labeled remodeler (green) and Hoechst-stained DNA (blue); the cross (yellow) indicates a representative location at which FCS measurements were performed. (**B**) Representative ACF curves for a remodeler subunit (gray) superimposed with the mean of normalized fits (red); residuals derived from individual fits of each ACF are shown on top. (**C**) Histograms of diffusion coefficients (*D*) of the fast (lighter colors) and slow (darker colors) fractions of BAF57 (top), BAF155 (middle) and BRG1 (bottom), as determined by fitting the ACF with a two-component model. (**D**) Histograms of anomalous diffusion coefficients (α) of the fast (lighter colors) and slow (darker colors) fractions of BAF57 (top), BAF155 (middle) and BRG1 (bottom), superimposed with a single-Gaussian fit (black) for the fast fraction and a double-Gaussian fit (black, with each Gaussian component in orange) for the slow fraction. *n* = 328, 398 and 447 independent measurements from 29, 42 and 48 cells for BAF57, BAF155 and BRG1, respectively for **C** and **D**.

**Table 1.** Diffusion parameters of key SWI/SNF remodeler subunits determined from intranuclear FCS measurements. Reported values are mean ± S.D. MWs include that of mEmerald.

| Subunit | MW (kDa) | $D$ ($\mu\text{m}^2\text{s}^{-1}$) | | Frequency $f$ | | $\alpha$ | | |
| --- | --- | --- | --- | --- | --- | --- | --- | --- |
|  |  | Fast | Slow | Fast | Slow | Fast | Slow,1 | Slow,2 |
| **BAF57** | 74 | $19.7 \pm 5.4$ | $0.42 \pm 0.67$ | $0.90 \pm 0.07$ | $0.10 \pm 0.07$ | $0.83 \pm 0.09$ | $0.67 \pm 0.06$ | $1.51 \pm 0.18$ |
| **BAF155** | 150 | $13.6 \pm 4.9$ | $0.47 \pm 0.64$ | $0.80 \pm 0.14$ | $0.20 \pm 0.14$ | $0.78 \pm 0.12$ | $0.58 \pm 0.02$ | $1.36 \pm 0.06$ |
| **BRG1** | 212 | $13.2 \pm 4.8$ | $0.29 \pm 0.34$ | $0.78 \pm 0.14$ | $0.22 \pm 0.14$ | $0.79 \pm 0.17$ | $0.87 \pm 0.05$ | $1.59 \pm 0.12$ |

Moreover, both fractions of each subunit exhibit anomalous diffusion (**Fig. 1D**). The fast fraction shows a narrow distribution with a mean anomalous diffusion coefficient of α_fast_ = 0.78–0.83, typical of non-Brownian subdiffusive behavior as a consequence of the crowded environment in the nucleoplasm (24–26). The slow fraction, however, has a much broader distribution for the anomalous diffusion coefficient, which can be decomposed into two subfractions with mean values of α_slow,1_ = 0.58–0.87 and α_slow,2_ = 1.36–1.59. While the α_slow,1_ subfraction resembles the subdiffusive characteristics of the fast fraction, the superdiffusive behavior of the α_slow,2_ subfraction has been previously attributed to chromatin-associated directed motion of DNA-interacting proteins (16, 27, 28). This phenomenon was also observed in our subsequent SMT measurements (**Fig. S4E** and **F**), thereby corroborating the fact that the slow fraction we observed indeed included chromatin-binding of the complex-bound SWI/SNF remodeler subunits.

### Single-molecule tracking quantifies transient and stable chromatin-binding of SWI/SNF remodelers

Due to the inability of FCS to accurately distinguish between different modes of diffusion and chromatin-binding interactions embodied by the slow fraction, we resorted to SMT measurements of individual Halo-tagged remodeler subunits, each labeled with the JF_549_ dye (29) (**Fig. 2A**). We optimized the intranuclear expression level of each remodeler subunit to be comparable to their endogenous levels in order to avoid overexpression-induced artifacts, and only selected cells within a narrowly controlled range of expression level for SMT measurements to ensure consistency (**Fig. S3**). We first performed fast tracking at 5.5 ms per frame (**Movie S1**). Analysis of the observed trajectories with the “Spot-On” framework (30) yielded displacement distributions that can only be satisfactorily fitted with a three-state model (**Figs. 2B** and **S4**). In addition to a population corresponding to the diffusion of the unincorporated individual subunits (similar to the fast fraction in FCS measurements), we further resolved the broad slow FCS fraction into two distinct modes with mean diffusion coefficients in the ranges of 0.10–0.12 µm^2^s^-1^ and 0.54–0.59 µm^2^s^-1^, respectively (**Fig. 2D**), with the former likely corresponding to transient chromatin-binding and the latter to diffusion, as evidenced by mean squared displacement (MSD) analysis (**Fig. S4D**). Importantly, the distributions of the diffusion coefficient and frequency for both modes overlap excellently between the three remodeler subunits, suggesting that they both correspond to the fully assembled complex in which all three subunits have been incorporated, as opposed to partially assembled complexes in which the various subunits will likely exhibit different dynamic properties depending on whether each of them has been incorporated.

**Figure 2.**
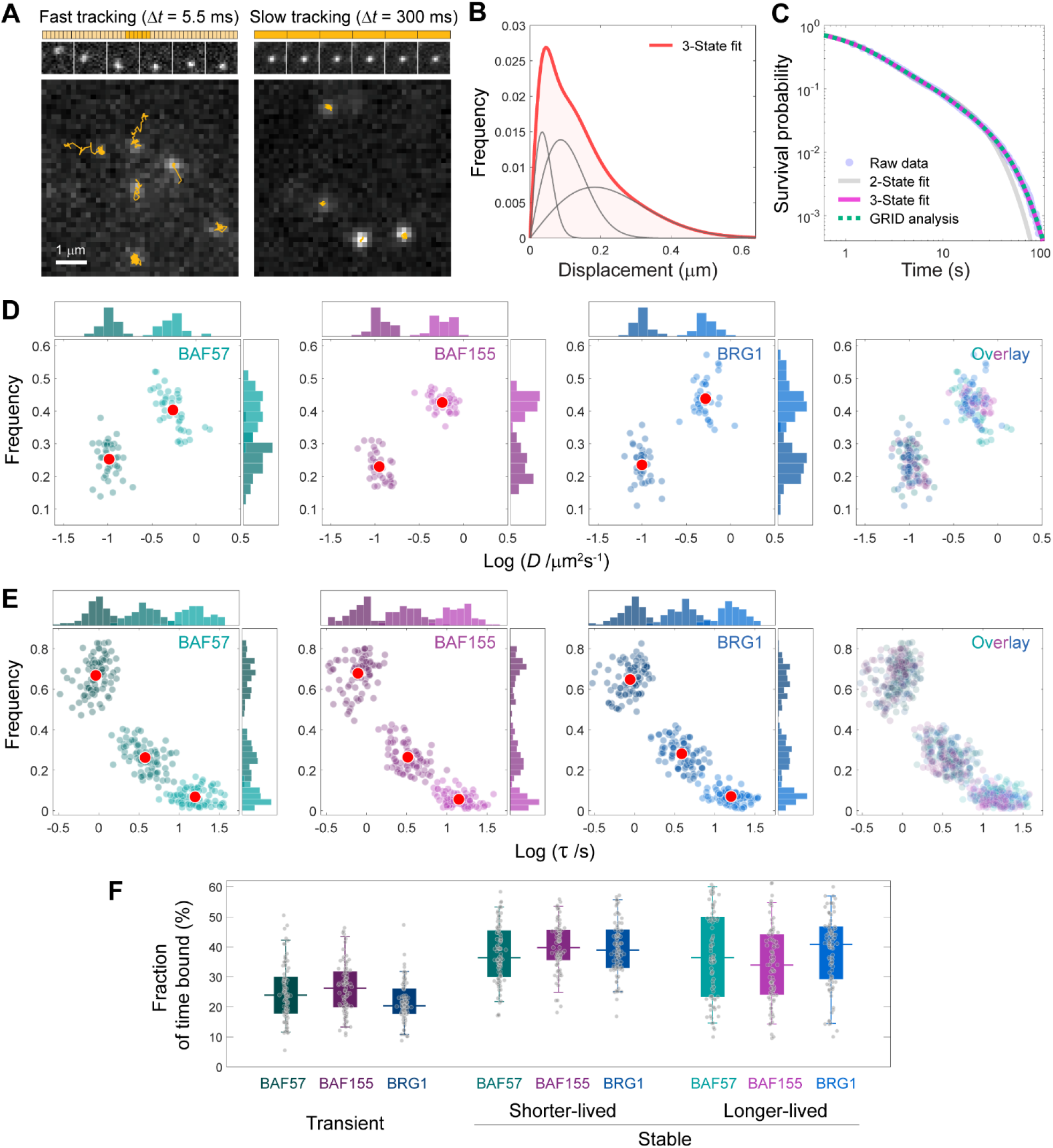
Live-cell single-molecule tracking quantifies diffusion and chromatin-binding of SWI/SNF remodelers. (**A**) Representative frames from SMT movies of a SWI/SNF remodeler subunit (BRG1 is shown here as example) in fast- (left, 5.5 ms per frame) or slow- (right, 300 ms per frame) tracking mode; individual trajectories (yellow) of diffusing (left) and chromatin-bound (right) remodeler molecules are superimposed on each frame. (**B**) A typical displacement histogram derived from fast-tracking trajectories in a single cell, satisfactorily fitted with a three-state model (red line). (**C**) A typical survival probability histogram derived from slow-tracking trajectories in a single cell, fitted with a two-state (gray) or three-state (pink) model or analyzed by GRID (dotted green). (**D**) Scatter plots of diffusion coefficient (*D*) and frequency associated with each molecular fraction resolved from fast-tracking trajectories for BAF57, BAF155, BRG1 and an overlay of the three plots. (**E**) Scatter plots of residence time (τ) and frequency associated with each molecular fraction resolved from slow-tracking trajectories for BAF57, BAF155, BRG1 and an overlay of the three plots. (**F**) Box-and-whisker plots of the relative fraction of time bound under each of the three modes resolved in **E** for BAF57, BAF155 and BRG1. In **D**–**F**, each dot denotes a single cell, and red circles in **D** and **E** denote mean values. *n* = 41 (BAF57), 38 (BAF155) and 39 (BRG1) cells for **D**, and 85 (BAF57), 81 (BAF155) and 90 (BRG1) cells for **E** and **F**.

In addition, we performed slow tracking (at 300 ms per frame) in order to blur out the rapidly diffusing remodeler molecules and facilitate the probing of binding events that take place on the second timescale (**Fig. 2A** and **Movie S2**). Given that most of the trajectories observed still consist of a mixture of diffusion and binding, we implemented a strategy to unambiguously discriminate binding from diffusion by scanning through all possible sub-trajectories and selecting only those that are spatially circumscribed within a confined area with an optimized radius (**Fig. S5A**). We also quantified the effect of photobleaching during acquisition to ensure that it minimally impacts the accuracy of our measurements (**Fig. S6**). The survival time distributions (31) computed from the binding trajectories were best fitted with a three-state model (**Fig. 2C**), consisting of a transient binding fraction with a mean residence time in the range of 0.83–0.97 s, as well as two stable binding fractions: a shorter-lived one with mean residence time in the range of 3.5–4.1 s, and a longer-lived one with mean residence time in the range of 15.2–17.3 s (**Fig. 2E** and **Table 2**). As validation, we analyzed the data with a genuine rate identification method (GRID), a recently developed analytical framework capable of robustly decomposing multi-component reaction systems in an unbiased fashion, which has been validated on DNA-binding proteins (32). The spectrum of residence times obtained by GRID showed three distinct peaks with residence times that agree excellently with those obtained from a three-state model (**Fig. 2C** and **Fig. S5B**), indicating that three distinct modes are indeed needed to fully account for the chromatin-binding dynamics observed. Finally, we computed the fraction of time the remodelers were bound under each mode (**Fig. 2F**), defined as the normalized product of the residence time and binding frequency associated with each mode (see **Materials and Methods** for details). We found that all three remodeler subunits exhibited similar fractions of time under each mode (with a mean fraction in the range of 22–26% for transient binding, 37–40% for shorter-lived stable binding and 34–39% for longer-lived stable binding). This, together with the fact that all three subunits showed similarly distributed residence times and binding frequencies (**Fig. 2E**), again confirms that we are indeed detecting chromatin-binding of the fully assembled remodeler complex.

**Table 2.** Chromatin-binding parameters of key SWI/SNF remodeler subunits determined from intranuclear SMT measurements. Reported values are mean ± S.D.

| Subunit | Residence time $\tau$ (s) | | | Frequency $f$ | | | Fraction of time bound $F$ | | |
| --- | --- | --- | --- | --- | --- | --- | --- | --- | --- |
|  | Transient binding | Stable binding |  | Transient binding | Stable binding |  | Transient binding | Stable binding |  |
|  |  | Shorter-lived | Longer-lived |  | Shorter-lived | Longer-lived |  | Shorter-lived | Longer-lived |
| **BAF57** | $0.97 \pm 0.29$ | $4.1 \pm 1.7$ | $17.3 \pm 7.6$ | $0.67 \pm 0.09$ | $0.26 \pm 0.07$ | $0.07 \pm 0.04$ | $0.25 \pm 0.09$ | $0.37 \pm 0.10$ | $0.38 \pm 0.15$ |
| **BAF155** | $0.83 \pm 0.29$ | $3.5 \pm 1.5$ | $15.2 \pm 6.2$ | $0.68 \pm 0.09$ | $0.26 \pm 0.07$ | $0.06 \pm 0.04$ | $0.26 \pm 0.09$ | $0.40 \pm 0.09$ | $0.34 \pm 0.13$ |
| **BRG1** | $0.93 \pm 0.30$ | $4.1 \pm 1.5$ | $17.1 \pm 6.6$ | $0.65 \pm 0.08$ | $0.28 \pm 0.07$ | $0.07 \pm 0.03$ | $0.22 \pm 0.07$ | $0.40 \pm 0.10$ | $0.39 \pm 0.12$ |

### DNA accessibility-dependent enhancement in remodeler binding dynamics is modulated by the bromodomain

We next probed how DNA accessibility could modulate the chromatin-binding dynamics of SWI/SNF remodelers in a quantitative manner by treating the cells with trichostatin A (TSA), a histone deacetylase inhibitor known to decondense chromatin and enhance DNA accessibility by promoting histone hyperacetylation (33). We found that enhancing DNA accessibility with TSA decreased the residence times (by an average of 16–21% for shorter-lived and 23–30% for longer-lived) and increased the frequencies (by an average of 14–30% for shorter-lived and 33–68% for longer-lived) of the stable binding modes for all three remodeler subunits (**Fig. 3A** and **B**). This is in line with our expectation that the more accessible chromatin requires less time to be remodeled at individual loci (which most likely takes place during stable binding events), and thereby allows the remodeler to remodel a larger number of loci per unit time. The residence time for transient binding, however, was less affected compared to that for stable binding, likely because this mode involves fast and nonspecific sampling of potential binding sites on chromatin, and the time required for each sampling event is less dependent on DNA accessibility. At the same time, since enhanced DNA accessibility enables the remodeler to engage in stable binding with a higher frequency, the frequency for transient binding is therefore reduced. Moreover, enhancing DNA accessibility does not significantly alter the fraction of time the remodeler stayed bound under each mode (**Fig. 3C**), thereby suggesting that DNA accessibility modulates remodeler–chromatin interactions by making binding more dynamic, but without shifting the overall partitioning of remodeler molecules among the different bound states.

**Figure 3.**
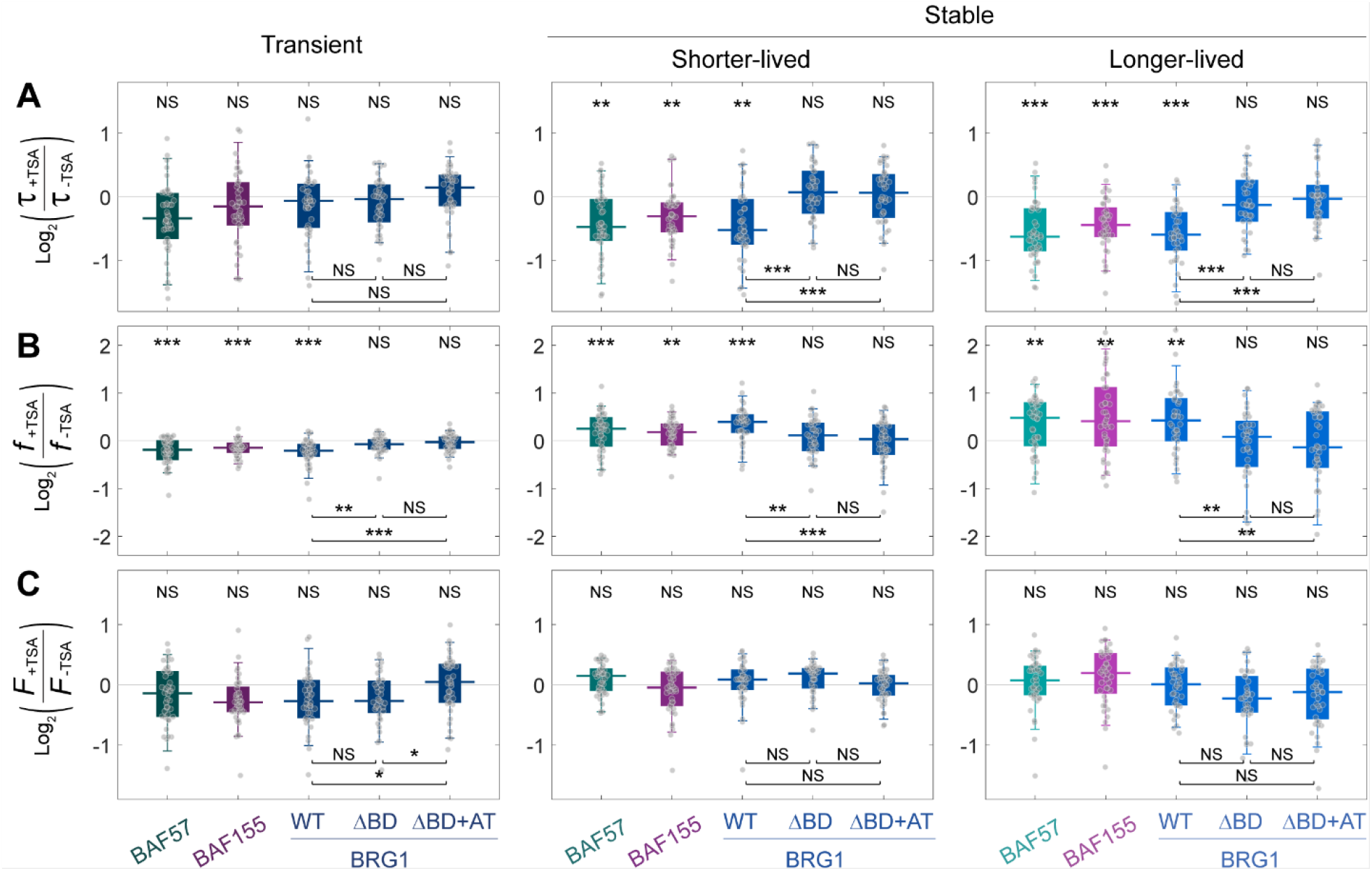
DNA-accessibility-dependent enhancement of SWI/SNF remodeler binding dynamics is modulated by the bromodomain. Box-and-whisker plots of fold changes (on a log_2_ scale) upon TSA treatment in (**A**) residence time (τ), (**B**) binding frequency (*f*) and (**C**) fraction of time bound (*F*) associated with each mode (transient, shorter-lived stable and longer-lived stable binding) for BAF57, BAF155 and BRG1 (both wildtype and two mutants with either BD or BD plus AT-hook truncated). Each dot denotes a single cell. *n* = 42 (BAF57), 40 (BAF155), 38 (BRG1), 36 (BRG1_ΔBD_) and 40 (BRG1_ΔBD+AT_) cells.

Furthermore, the dynamic targeting and binding of the BAF180 subunit unique to the PBAF remodeler subtype has recently been shown to be regulated by the bromodomain (BD) (19). Among the three SWI/SNF remodeler subunits under our examination, BRG1 harbors a BD that enables it to bind to the major DNA groove and recognize acetylated lysines on histone tails (34–36). Hence, in order to probe the generic role of BD in SWI/SNF remodeler dynamics and its dependence on DNA accessibility, we compared the chromatin-binding of two naturally existing mutants of BRG1 that truncates either the BD (E1449X (37), thereafter denoted as ΔBD) or the BD plus the preceding AT-hook (a short arginine/lysine-rich motif that binds to the minor DNA groove formed by AT-rich DNA elements (38)) (R1415X (39), thereafter denoted as ΔBD+AT) with wildtype BRG1 (**Fig. 3A** and **B**, and **Fig. S7**). For both mutants, we found that deleting the BD abrogates the enhanced binding dynamics induced by TSA, indicating that BD plays a generic role in modulating the binding dynamics of SWI/SNF remodelers in a DNA-accessibility-dependent manner. However, the fact that deleting the AT-hook did not further contribute to this effect suggests that the AT-hook is not involved in binding acetylated histone tails.

### Super-resolved density mapping of SMT trajectories reveals intranuclear remodeler binding “hotspots”

To furnish spatial contexts to the intranuclear dynamics observed above, we reasoned that the density of displacements (or steps) detected in SMT trajectories at a particular intranuclear location serve as a measure of the frequency with which that location is visited or resided upon by the remodeler. Hence, we devised a strategy, termed STep Accumulation Reconstruction (STAR), to map the distribution of local displacement density in SMT binding trajectories in a similar way as localization coordinates are used in the reconstruction of superresolution microscopy images (**Fig. S8**). While our approach is conceptually akin to methods such as sptPALM (40) and other variants (41–44), STAR substantially expands the scope and utility of previous methods that focus only on diffusion by also enabling the mapping of binding events density across the cell nucleus. The resulting super-resolved density maps for each of the three binding modes (see **Materials and Methods** for details) revealed distinctly and heterogeneously distributed clusters across the nucleoplasm, with each cluster corresponding to a binding “hotspot” where the remodeler binds preferentially and repeatedly; in contrast, the corresponding density map for diffusion exhibited a more homogeneous distribution (**Fig. 4A**). Ripley’s K-function analysis indicated that these hotspots have a characteristic radius of *r*_c_ ∼140 nm (regardless of whether all binding modes or only stable binding modes are considered, **Fig. 4B**), and consist of an average of 6.2 binding events per cluster (**Fig. 4C**). Furthermore, to probe the temporal correlation between binding events within a hotspot, we compared the probabilities of observing two consecutive (P(A∩B)) vs. two independent (P(A)·P(B)) binding events (**Fig. 4D**). Interestingly, the probability of observing two consecutive longer-lived stable binding events within a cluster is significantly higher than that of observing two independent such events, as compared to other sequential combinations of longer-lived and shorter-lived binding events (all of which are largely independent of each other). The fact that a longer-lived binding event is more likely to be followed by another longer-lived binding event suggests that the genomic binding sites within these hotspots could be predisposed towards or induced upon longer-lived binding to favor further such binding, as a potential strategy to promote remodeling at these sites. Expectedly, most of the hotspots disappeared in the presence of PFI-3 (a BRG1 BD inhibitor (45)) or when tracking only the HaloTag (which does not bind to DNA) (**Fig. S9**), thereby validating the veracity of the binding hotspots we observed.

**Figure 4.**
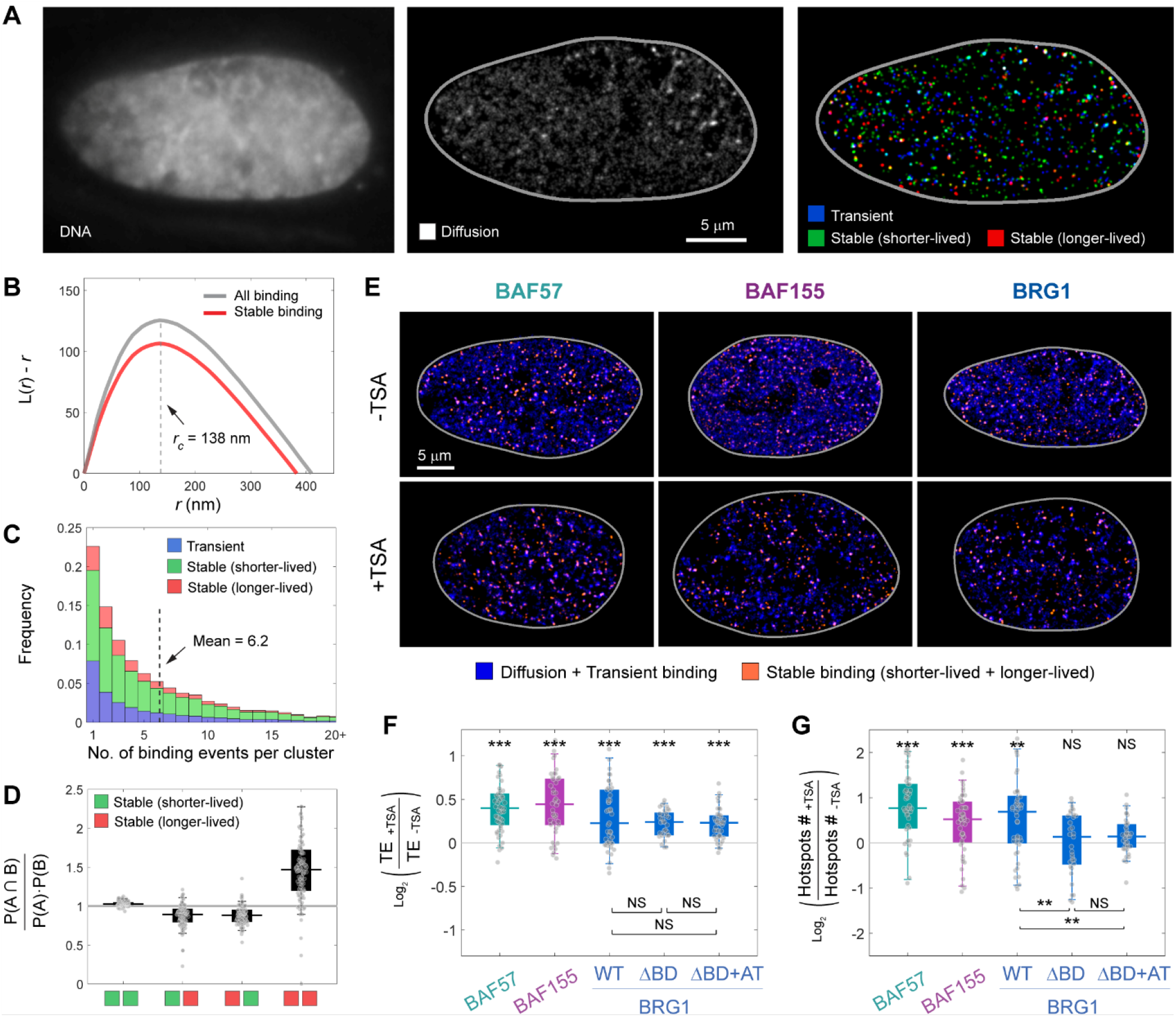
Super-resolved STAR mapping reveals nanoscale remodeler binding hotspots that exhibit clustered organization of binding events. (**A**) A live HeLa cell nucleus showing Hoechst-stained DNA background (left) and the corresponding STAR maps for diffusion (middle) and binding (right, color-coded according to binding mode). Gray line delineates nuclear boundary. (**B**) Plots of Ripley’s K-function for either all binding events (gray) or stable binding events only (red) reveal the existence of nanoscale clusters corresponding to remodeler binding hotspots; dotted line indicates typical cluster radius (*r*_c_) corresponding to the peak of the K-function. (**C**) Histogram of the number of binding events per cluster, color-coded according to binding mode; dotted line indicates mean value. (**D**) Box-and-whisker plot of the ratio between the probability of observing two consecutive binding events and that of observing two independent binding events in a cluster, 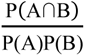 (where A and B denotes either a shorter-lived (green) or a longer-lived (red) stable binding event), indicates that consecutive longer-lived binding events are statistically more correlated as compared to all other sequential combinations (which are largely independent of each other). (**E**) STAR maps for diffusion and transient binding combined (deep blue) and stable binding (both shorter-lived and longer-lived, light red) for each of the three remodeler subunits before (top row) and after (bottom row) TSA treatment. (**F** and **G**) Box-and-whisker plots of fold changes (on a log_2_ scale) upon TSA treatment in (**F**) targeting efficiency (TE) and (**G**) number of stable binding hotspots for BAF57, BAF155 and BRG1 (both wildtype and two mutants with either BD or BD plus AT-hook truncated). In **D**, **F** and **G**, each dot denotes a single cell. *n* = 107 (BAF57) cells for **D**, and 44 (BAF57), 42 (BAF155), 45 (BRG1), 33 (BRG1_ΔBD_) and 37 (BRG1_ΔBD+AT_) cells for **F** and **G**.

Moreover, upon enhancing DNA accessibility with TSA treatment, the binding maps for all three remodeler subunits exhibited a marked reduction in the density of diffusion and transient binding events relative to the stable binding events (**Fig. 4E**). To better quantify this trend, we defined the targeting efficiency (TE) of each remodeler as the ratio between the intranuclear space explored by stable binding versus that by diffusion and transient binding combined (see **Materials and Methods** for details), which provides a convenient metric for how efficiently each remodeler can specifically target the desired genomic sites for stable binding. As expected, TSA treatment significantly enhanced the TE for all three subunits (**Fig. 4F**) and increased the number of stable binding hotspots detected (**Fig. 4G**), thereby suggesting that enhancing DNA accessibility can either expose more genomic spaces for remodelers to bind or generate new binding hotspots by virtue of the higher frequency for sampling genomic target sites as a result of enhanced binding dynamics. However, deleting the BD in BRG1 using the two aforementioned mutants (ΔBD and ΔBD+AT) did not abrogate this enhanced TE (**Fig. 4F**), thereby suggesting that BD does not provide additional targeting capability to the hyperacetylated and more accessible chromatin upon TSA treatment. On the other hand, deleting the BD did abrogate the increased number of stable binding hotspots induced by TSA (**Fig. 4G**), indicating that the additional binding hotspots are a consequence of the enhanced binding dynamics (particularly the higher binding frequency, **Fig. 3B**), as opposed to the expanded genomic spaces that can be sampled by the remodelers upon TSA treatment.

### Distinct, multi-modal alterations in DNA-binding dynamics are associated with various cancer-implicated BRG1 mutants

Despite the fact that BRG1 is one of the most frequently mutated SWI/SNF remodeler subunits in human cancers (13), how these mutations impact the DNA-binding dynamics of the remodeler complex in relation to various cancers remains poorly understood. To address this key deficiency, we examined seven common BRG1 mutants implicated in a variety of cancers across tumor types, including those that affect the skin, lung, liver, colon, bladder, kidney, ovary and placenta (**Table 3**); their respective locations are mapped onto the domain organization and structure of BRG1 in complex with a nucleosome (**Fig. 5A** and **B**, and **Fig. S10**) (6). Among the three point mutations, E861K (46) is located within the central ATPase domain, which consists of two conserved RecA-like lobes known as DExx and HELICc domains, and is the catalytically active domain involved in ATP binding/hydrolysis and DNA translocation during remodeling (47–49). In addition, P1456L occurs in the linker between the AT-hook and BD, while R1502H occurs within the BD (34). Among the four truncation mutants, Δ530–807 (50) has parts of the ATPase domain and the HSA domain (which regulates remodeling activity in tandem with its neighboring post-HSA domain (9)) as well as the entire BRK domain (which consist of a potential protein-binding motif with unknown function (51)) deleted, while Δ851–1138 (50) has a part of the ATPase domain deleted. Finally, Q1304X (52) harbors a stop codon mutation that leads to the C-terminal truncation of the BD, AT-hook and the SnAC/post-SnAC domains (which regulate ATPase activity and nucleosome mobilization (53, 54) and facilitate remodeler anchoring to the nucleosome core during remodeling (6)), while Q164X harbors a stop codon mutation that leads to the truncation of all functional domains in BRG1 (50).

**Figure 5.**
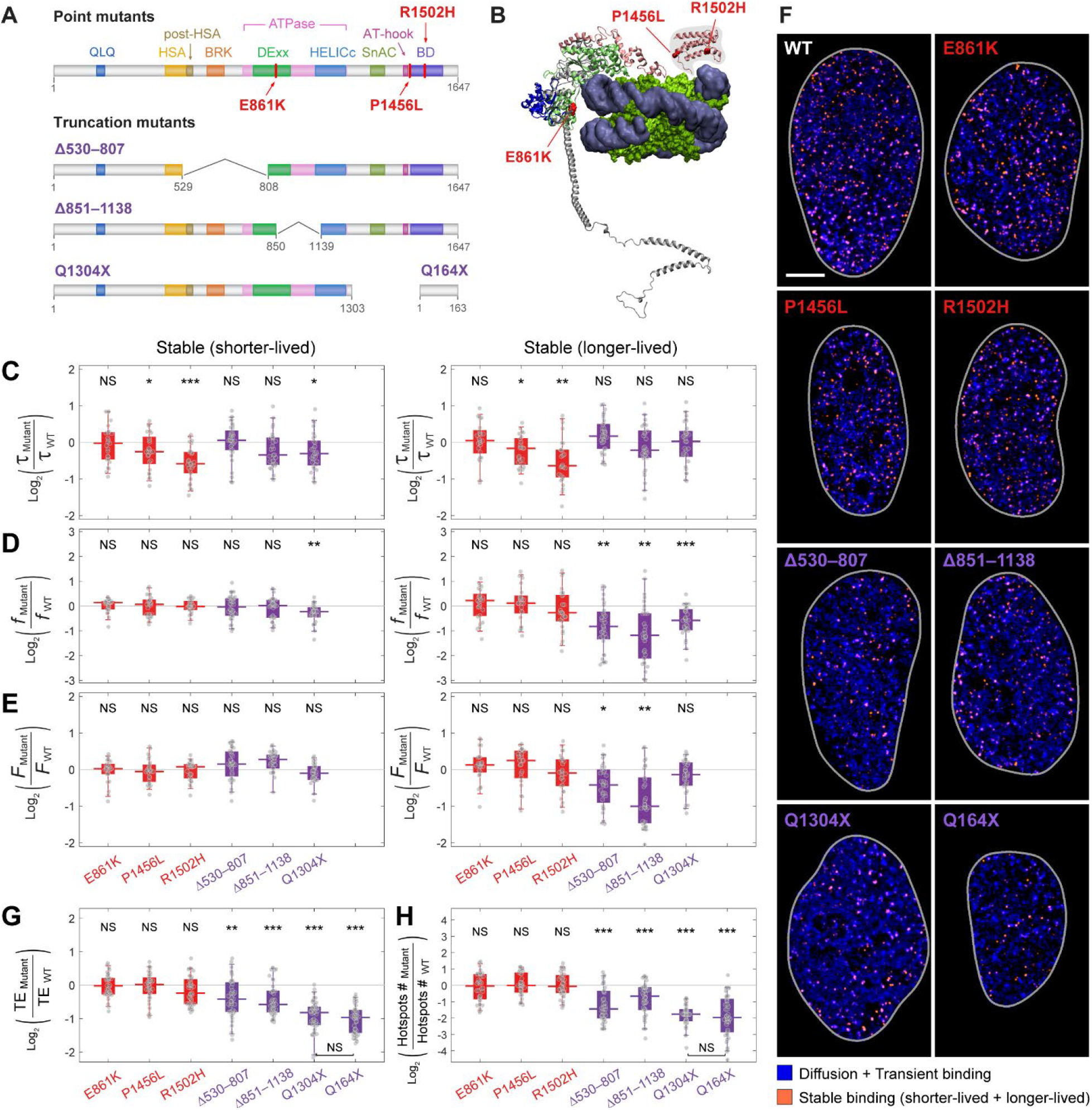
BRG1 mutants implicated in diverse cancers exhibit distinct and multi-modal alterations in DNA-binding dynamics. (**A**) Overview of domain organization of BRG1 and the locations of seven point/truncation mutants examined. (**B**) Cryo-EM structure of BRG1 (gray, in cartoon representation) in complex with nucleosome (blue-gray and yellow-green, in surface representation) (6). Point mutations E861K, P1456L and R1502H are shown in red, and truncation in mutants Δ530–807, Δ851–1138 and Q1304X are shown in blue, green and pink, respectively. Residues 1–335 (which is largely intrinsically disordered and contains the truncation mutant Q164X), 532–679 (part of the truncation in mutant Δ530–807) and 1419–1647 (containing the BD), however, were not resolved in the original structure. The structure of the BD (pink) was separately resolved in complex with DNA (34), and is placed here (shaded area) to illustrate its approximate orientation in the overall complex. (**C** to **E**) Box-and-whisker plots of fold changes (on a log_2_ scale) in (**C**) residence time (τ), (**D**) binding frequency (*f*) and (**E**) fraction of time bound (*F*) associated with shorter-lived and longer-lived stable binding for each of the seven mutants (except Q164X), relative to that for wildtype BRG1. (**F**) STAR maps for diffusion and transient binding combined (deep blue) and stable binding (both shorter-lived and longer-lived, light red) for each of the seven mutants, together with wildtype BRG1 as reference. Gray line delineates nuclear boundary. (**G** and **H**) Box-and-whisker plots of fold changes (on a log_2_ scale) in (**G**) targeting efficiency (TE) and (**H**) number of stable binding hotspots for each of the seven mutants, relative to that for wildtype BRG1. In **C**–**E**, **G** and **H**, each dot denotes a single cell. *n* = 25 (E861K), 25 (P1456L), 26 (R1502H), 32 (Δ530–807), 30 (Δ851–1138) and 29 (Q1304X) cells for **C** to **E**; and 34 (E861K), 34 (P1456L), 35 (R1502H), 39 (Δ530–807), 38 (Δ851–1138), 41 (Q1304X) and 39 (Q164X) cells for **G** and **H**.

**Table 3.**
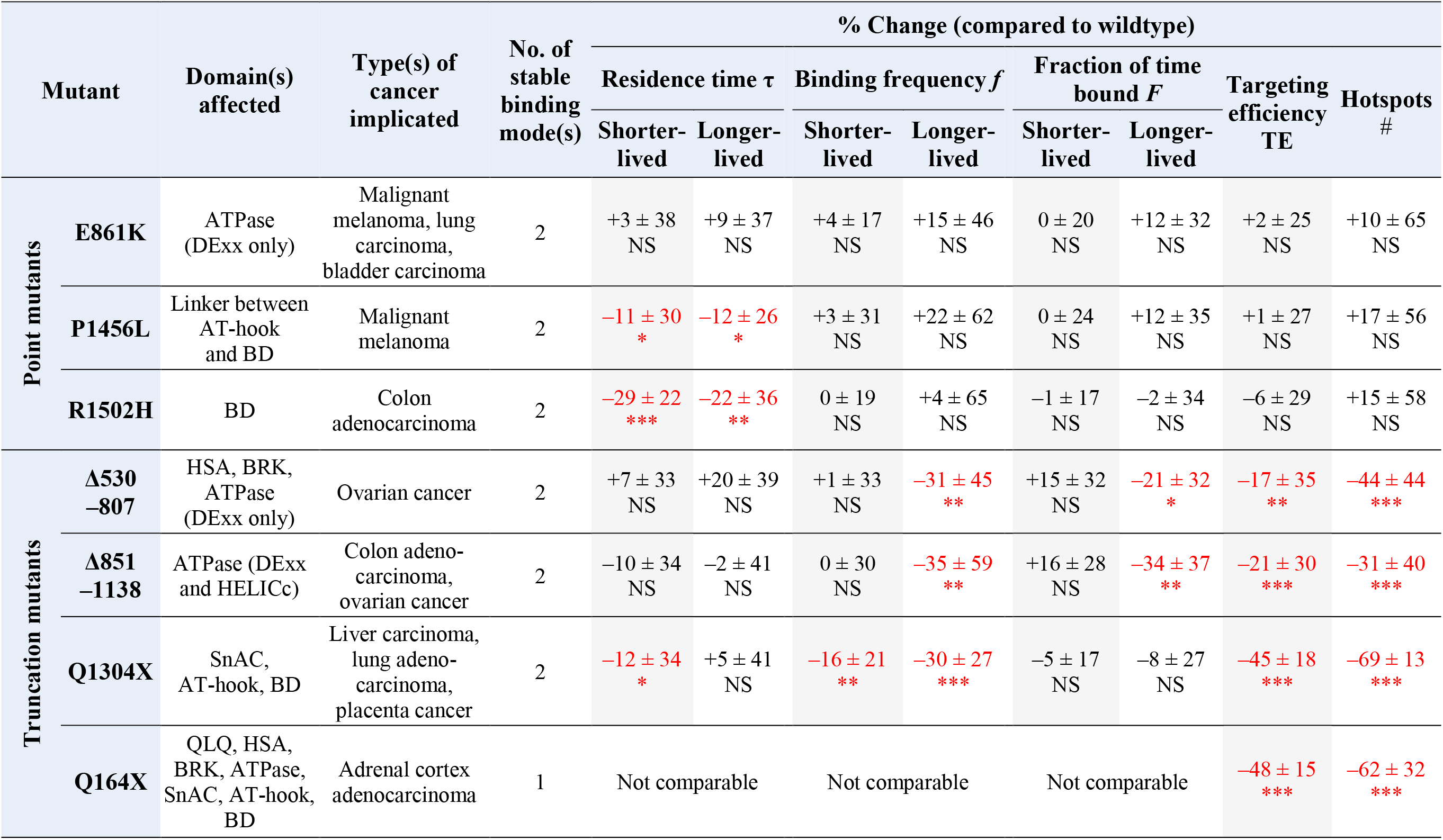
Mutant-specific signatures in chromatin-binding dynamics associated with seven cancer-implicated BRG1 mutants. Reported values are mean ± S.D. “+” and “–” denotes increase and decrease in value, respectively. The values for τ, *f* and *F* for mutant Q164X (which shows only one stable mode) are not comparable to those of other mutants (which show two distinct stable modes).

We first validated that all mutants were expressed at comparable levels and exhibited similar intranuclear localization patterns as wildtype BRG1 (**Fig. S11**). Performing SMT measurements, quantifications and STAR mapping as described above (**Fig. 5C to H** and **Table 3**), we found that the point mutation R1502H in the BD resulted in significantly reduced residence times for both modes of stable binding, possibly as a result of replacing the positively charged arginine finger that interacts with the negatively charged DNA backbone with the more neutral and shorter histidine residue, thereby destabilizing remodeler–DNA binding. Similarly, replacing the proline residue in the linker between the AT-hook and BD with leucine (P1456L) partially disrupts the conformation of the linker that positions both domains into proper contact with DNA (55), thereby mildly reducing the residence times for stable binding. Finally, point mutation E861K exhibited no significant change in any of the parameters measured, in line with the fact that it does not lie within any conserved motif (involved in either ATP- or DNA-binding) of the DExx domain (49).

In contrast to the point mutants, the C-terminal truncation mutant Q1304X exhibited significant reductions in the frequencies of both stable binding modes, targeting efficiency and number of stable binding hotspots. This is in agreement with the roles of BD in binding DNA and acetylated histone tails as well as of the recently postulated post-SnAC domain in binding the nucleosome core and facilitating its anchoring during remodeling (6). The impaired ability of this mutant to bind nucleosome, as manifested by the drop in stable binding frequencies, increases the time spent in diffusing and nonspecifically sampling the genomic space, hence leading to reductions in TE and number of binding hotspots. Moreover, truncation mutants Δ530–807 and Δ851–1138 also showed significant reductions in both the frequency as well as the fraction of time bound under the longer-lived stable mode, targeting efficiency and number of stable binding hotspots. This can be similarly attributed to the fact that both mutants involve truncation of part of the ATPase domain, which contains eight conserved motifs (**Fig. S10**) variously involved in ATP-binding, DNA-binding and intramolecular interactions: the truncated region of Δ530–807 harbors the Q-motif and motif I (involved in ATP-binding/hydrolysis), while the truncated region of Δ851–1138 harbors motifs II, III and IV (involved in ATP-binding/hydrolysis, intramolecular interactions and DNA-binding) (49). In addition, the post-HSA domain truncated in Δ530–807 is known to also interact with DNA and contribute to the overall affinity of the remodeler complex (56) as well as regulating the ATPase domain by mechanically connecting the ATP binding site with the HSA domain (9). Moreover, the proper orientations and positioning of all domains C-terminal to the truncated region (especially the SnAC/post-SnAC domains and BD) in relation to the nucleosome are likely disturbed in both mutants, potentially leading to misalignment or even conformations that could sterically hinder the remodeler from establishing stable contacts with the nucleosome. These effects combined account for the impaired ability of both mutants not only to bind DNA but also to engage in ATP hydrolysis-driven remodeling (which likely takes place during longer-lived binding). The resulting drop in the frequency of longer-lived stable binding also translates into reduction in the fraction of time bound under this mode (**Fig. 5E**), hence shifting the molecular partitioning of the remodeler among the various modes.

Finally, mutant Q164X exhibited not only drastically reduced chromatin-binding, targeting efficiency and number of stable binding hotspots similar to the other three truncation mutants, but also showed only a single stable binding mode (with a residence time of 6.4 ± 1.6 s) that is distinct from the two stable binding modes for wildtype BRG1 (with residence times of 4.1 ± 1.5 s and 17.1 ± 6.6 s, respectively). This was further confirmed by FCS measurement, in which the contribution of the slow fraction was found to be drastically reduced (**Fig. S12**). In particular, the marked (close to 3-fold) reduction in the number of stable binding hotspots is in line with the previous finding that removing the BRG1/BRM subunit reduces the chromatin affinity of mammalian SWI/SNF complexes to a residual level (57). As such, the reduced chromatin-binding/targeting we observed could be a consequence of such residual affinity exhibited by the remodeler complex that has incorporated this mutant. Even though binding can still be achieved (albeit at a much lower level) via this single stable mode through the interactions of other remodeler subunits with the nucleosome, no subsequent remodeling can be carried out by this mutant due to the deletion of all functional domains.

Collectively, these findings revealed a multi-modal spatio-temporal landscape used by SWI/SNF remodelers via the dynamic modulation of the efficiency of targeting/binding chromatin as well as the differential partitioning of the remodeler molecules among various modes to regulate remodeler–chromatin interactions *in vivo*.

## DISCUSSION

### Distinct modes of diffusion/chromatin-binding dynamics resolved for SWI/SNF remodelers

Despite our knowledge about the biochemistry, structure and genetics of the human SWI/SNF chromatin remodeler subfamily, their intranuclear organization and dynamics in space and time remain much less understood, nor do we know how misregulation of such organizational dynamics might underpin the various types of cancer in which they are implicated. Here, we addressed these deficiencies by performing, to our knowledge, the first systematic single-molecule quantification of the live-cell dynamics of the fully assembled human SWI/SNF remodeler complex using two complementary approaches. We first resolved distinct modes of intranuclear dynamics for three SWI/SNF remodeler subunits using FCS, revealing a large (∼80%) fast fraction corresponding to the diffusion of unincorporated individual subunits, and a small (∼20%) but much broader slow fraction that corresponds to the diffusion and/or chromatin-binding of the assembled complex (both partially and fully). To further resolve this slow fraction, we resorted to SMT measurements, which revealed one transient binding and two stable binding modes for all three remodeler subunits. Critically, the fact that all three subunits exhibited similarly distributed residence times, binding frequencies and fractions of time bound confirms that the three binding modes we resolved indeed correspond to the binding of the fully assembled remodeler complex. Among them, the transient binding mode likely corresponds to nonspecific interaction or sampling events, as have been similarly observed for other DNA-binding proteins such as transcription factors (TFs) (59) and DNA replication factors (60). In contrast, the consistent observation of two distinct stable binding modes for all three remodeler subunits can be potentially accounted for by the fact that different subtypes of the SWI/SNF complexes tend to engage different chromatin sites: it is known that BAF remodelers primarily target distal enhancer regions whereas PBAF remodelers primarily target promoter regions (61, 62). The varying accessibility of the genome could also play a role, as PBAF has been found to display shorter residence times at the more open and presumably easier-to-remodel euchromatin, as compared to the denser and harder-to-remodel heterochromatin (19). Furthermore, different types of remodeling activities (*e.g.* nucleosome translocation vs. eviction), which involve complex multi-step pathways (4), will also likely require different binding durations to complete. Finally, the anomalous diffusion behaviors (both subdiffusive and superdiffusive) observed by both FCS and SMT attest to the complex dynamics present within the highly crowded intranuclear environment, in the form of either restricted motion due to the dense chromatin meshwork or directed motion along with chromatin for DNA-binding proteins (58).

Our findings are in general agreement with a FCS-based model previously proposed for the human ISWI remodeler subfamily, in which the majority of the remodeler complexes continuously sample the nucleosomes via transient binding while a small fraction stably binds to chromatin for extended durations to carry out remodeling (16), although our SMT measurements have enabled us to further resolve the stable binding fraction of the SWI/SNF subfamily with much greater detail. In addition, a recent study on six different yeast remodeler subfamilies (including SWI/SNF) also uncovered only one stable binding mode for all remodelers (with a mean residence time of 4.4 s for the SWI/SNF subfamily) (18), as opposed to the two distinct stable modes we found for human SWI/SNF remodelers. This difference could potentially be reconciled by the fact that the yeast genome is less complex in architecture, with much of the genome being constitutively open for transcription (63), and hence requires less sophisticated mechanisms to remodel.

### Intranuclear hotspots mapping sheds insight into the spatio-temporal organization of selective remodeler engagement with genomic target sites

Adding onto the temporal aspects of remodeler binding dynamics, the STAR mapping strategy we devised offers a powerful way for directly visualizing the spatial landscape of remodeler binding in a mode-specific manner. With this capability, we revealed numerous nanoscale and heterogeneously distributed binding hotspots across the nucleoplasm for SWI/SNF remodeler complexes, in which multiple binding events (for both transient and stable binding) preferentially cluster. The typical size we observed for these hotspots (∼140 nm in radius) agrees excellently with that previously found for BAF180 hubs (∼250 nm in diameter) (19). More importantly, the enrichment of sequential longer-lived stable binding events in these hotspots, revealed through statistical dependency tests (**Fig. 4D**), suggests that they likely correspond to genomic loci where the target sites are predisposed towards or induced upon longer-lived binding to promote remodeling at these sites, as a potential strategy used by SWI/SNF remodelers to selectively engage genomic target sites to provide DNA access for other chromatin-dependent processes. Similar hotspots have also been previously observed for ISWI remodelers; these hotspots are bound by the remodelers for durations similar to those we observed for SWI/SNF remodelers (both in the range of seconds to minutes), and were found to be enriched at DNA damage repair sites and replication foci in S phase cells (16).

### Roles of BD in modulating DNA-accessibility-dependent remodeler binding/targeting to chromatin

Our finding that enhancing DNA accessibility makes remodeler binding more dynamic (as manifested by shorter residence times and higher frequency for both modes of stable binding) is consistent with the enhanced targeting efficiency observed upon TSA treatment, since the remodeler can now better access the chromatin and engage in stable binding more frequently (as opposed to diffusing in the nucleoplasm or nonspecifically sampling the genome). Moreover, the fact that deleting the BD in BRG1 abrogates such enhanced binding dynamics points to a generic role played by BD in modulating the chromatin binding of SWI/SNF remodelers in a DNA-accessibility-dependent manner. However, deleting the BD in BRG1 does not impact the enhanced targeting efficiency associated with TSA treatment, thereby suggesting that the previously proposed role of BD in targeting the hyperacetylated and more accessible chromatin (64) might need to be reconsidered. Moreover, the fact that deleting the BD in BRG1 does abrogate the higher number of stable binding hotspots associated with TSA treatment indicates that enhanced binding dynamics plays a more prominent role in the generation of new binding hotspots as compared to the additional genomic space that is exposed to the remodelers upon enhancing DNA accessibility. Finally, deleting the AT-hook does not further contribute to all of these effects, in line with the previous finding that the AT-hook is not involved in binding acetylated histone tails (34).

### A structure-dynamics framework for elucidating cancer-mutant-specific alterations in remodeler–chromatin interactions

The multi-modal changes in chromatin-binding dynamics associated with various cancer-implicated mutants revealed by our single-molecule measurements can be rationalized with a framework that integrates various structural features of BRG1 in the context of nucleosome binding (**Fig. 5** and **Fig. S10**). Among the mutants examined, the reduction in the residence times for both shorter- and longer-lived stable binding exhibited by point mutants R1502H and P1456L is in line with previous *in vitro* binding affinity measurements and NMR chemical shift perturbations observed for these mutants (34). In contrast, truncation mutants Δ530–807, Δ851–1138 and Q1304X are all characterized by a significantly reduced frequency of longer-lived stable binding, indicating a lowered probability of engaging in binding that will likely lead to remodeling, since binding frequency provides a metric for the “on”-rate of remodeler binding (32). However, given that the residence times for these mutants remain largely unchanged, the mutant remodeler complexes, upon successful engagement, can nonetheless carry out remodeling in a normal manner similar to wildtype BRG1. Moreover, the fact that the molecular partitioning (a metric for binding affinity) of the longer-lived stable mode is shifted for the two internal truncation mutants (Δ530–807 and Δ851–1138) but not for the C-terminal truncation mutant (Q1304X) points to the importance of proper alignment between key remodeler domains (especially the SnAC/post-SnAC and BD) and structural features of the nucleosome. Finally, further truncation beyond the SnAC domain (Q164X) does not lead to further reduction in targeting efficiency and number of stable binding hotspots (**Fig. 5G** and **H**), hence suggesting that the SnAC domain, AT-hook and BD may collectively constitute a sufficient functional module in BRG1 that synergistically modulates the targeting, positioning and binding of the remodeler complex to nucleosome. However, the observation of only a single stable binding mode with a residence time closer to that of the shorter-lived stable mode indicates that the binding of this mutant will most likely not lead to any remodeling, and illustrates the diverse range of dynamic parameters that could be impacted as a result of cancer-associated remodeler mutations. Finally, point mutant E861K does not exhibit a significant change in any of the parameters measured, contrary to previous FRAP measurements which found faster recovery after photobleaching compared to wildtype BRG1 (46). However, the recovery observed here occurs on a minute timescale and corresponds to slower changes in remodeler**–**chromatin interactions than that probed by our much more localized single-molecule measurements on a second timescale. Similarly, two other point mutations T910M and R1192C located in the DExx and HELICc domain, respectively, also do not affect the residence times, binding frequencies or fractions of time bound for BRG1 (**Fig. S13**), likely because these residues are involved in ATP binding/hydrolysis and not in nucleosome-binding directly.

Overall, the approaches developed in our study have revealed critical intranuclear dynamics of the SWI/SNF remodeler subfamily, and paved the way for further investigations into the organization and dynamics of chromatin remodeling. In particular, our findings establish the biophysical basis for aberrant remodeler–chromatin interactions associated with seven SWI/SNF remodeler mutants implicated in various cancers, and could potentially serve as a unique set of identifying yardsticks for these mutants. More fundamentally, they also revealed a much broader and multi-modal landscape at work to regulate remodeler dynamics beyond that captured by residence time and binding fraction, and argue for the adoption of other equally revealing but often under-explored parameters in both space and time in order to comprehensively describe remodeler–chromatin interactions *in vivo*. However, the lack of observable change in the three aforementioned point mutants underscores the fact that even such a wide coverage of parameter space as ours still cannot fully capture the consequences of cancer-causing remodeler mutations, and calls for a holistic approach to reveal their functional impact (*e.g.* by simultaneously monitoring related processes such as TF binding, transcription, remodeling activity, *etc.*). Correlating remodeler binding hotspots with other key nuclear structures (*e.g.* transcription factories (65), DNA replication foci (66), DNA loops (67) and epigenetic signatures) will also enable us to pinpoint the molecular nature and chromatin microenvironments of these hotspots. Finally, quantifying the impact of multiple mutations that co-exist in the same cancer cell could potentially reveal synergistic effects not manifested by individual remodeler mutants, thereby shedding light on the mechanistic consequences of remodeling misregulation in more disease-relevant contexts.

## MATERIALS AND METHODS

### Reagents

All restriction enzymes, NEBuilder HIFI DNA Assembly Master Mix (E2621L), KLD Enzyme Mix (M0554S) and the Quick Ligation Kit (M2200L) were purchased from New England Biolabs (Ipswich, USA). All other chemicals were purchased from Merck (Darmstadt, Germany).

### Constructs generation

Human BAF57 and BAF155 gene fragments were amplified from plasmids pBS-hBAF57 (Addgene ID #17877) and pBS-hBAF155 (Addgene ID #17876), respectively, while the human BRG1 gene fragment (splicing isoform 2 (1614 aa), which differs from the canonical isoform 1 (1647 aa) by the deletion of 33 amino acids (aa 1259-1291) from exon 28, outside of any functional domains) was amplified from the plasmid pCS2-hBRG1 (a gift of Nicolas Plachta, University of Pennsylvania). C-terminal mEmerald fusion constructs (pBAF57-mEm, pBAF155-mEm and pBRG1-mEm) were generated by cloning each gene fragment into a pCS2-mEmerald backbone plasmid (a gift of Nicolas Plachta, University of Pennsylvania) via Gibson assembly using the NEBuilder HIFI DNA Assembly Master Mix according to manufacturer’s protocol.

C-terminal HaloTag fusion construct for BAF57 (pBAF57-HTC) was generated by amplifying the HaloTag backbone from the pHTC HaloTag® CMV-neo vector (G7711, Promega) and digesting it with AgeI and KpnI, followed by ligation with the BAF57 gene fragment using the Quick Ligation Kit. C-terminal HaloTag fusion construct for BAF155 (pBAF155-HTC) was generated by amplifying the HaloTag backbone from a pCS2-HaloTag vector (in-house), followed by Gibson assembly with the BAF155 gene fragment using the NEBuilder HIFI DNA Assembly Master Mix.

N-terminal HaloTag fusion construct for BRG1 (pHTN-BRG1) was generated by digesting the pHTN HaloTag® CMV-neo vector (G7721, Promega) with PvuI and XbaI, followed by ligation with the BRG1 gene fragment using the Quick Ligation Kit. N-terminal HaloTag fusion constructs for BRG1 point mutants (pHTN-BRG1_E861K_, pHTN-BRG1_T910M_, pHTN-BRG1_R1192C_, pHTN-BRG1_P1456L_ and pHTN-BRG1_R1502H_) were generated via site-directed mutagenesis by introducing each point mutation via the respective primer during pHTN-BRG1 amplification. The linear fragment obtained was then subjected to the KLD Enzyme Mix to re-circularize the plasmid. N-terminal HaloTag fusion constructs for BRG1 truncation mutants (pHTN-BRG1_Δ530–807_, pHTN-BRG1_Δ851–1138_, pHTN-BRG1_Q1304X_ and pHTN-BRG1_Q164X_, as well as the BD truncation mutants pHTN-BRG1_ΔBD_ and pHTN-BRG1_ΔBD+AT_) were generated by amplifying the desired pHTN-BRG1 sequence while omitting the respective truncated region. The linear fragment obtained was then subjected to the KLD Enzyme Mix to re-circularize the plasmid. N-terminal mEmerald fusion construct for the BRG1_Q164X_ mutant (pmEm-BRG1_Q164X_) was generated by amplifying the desired sequence from a pCS2-mEm-BRG1 plasmid (in-house) while omitting the truncated region. The linear fragment obtained was then subjected to the KLD Enzyme Mix to re-circularize the plasmid. The location of each mutation was mapped to the corresponding position in BRG1 isoform 1 (UniProt entry: P51532.1). The proper generation of all constructs generated was confirmed by sequencing. All primers used to generate the constructs are listed in **Table S1**.

### Cell culture and transfection

Immortalized HeLa cell line (ATCC CCL-2, a gift from Thorsten Wohland, National University of Singapore) was cultured in a 25-cm^2^ flask using HyClone Dulbecco’s modified Eagle’s medium (DMEM) with high-glucose (SH30022.01, Cytiva) supplemented with 10% (v/v) fetal bovine serum (FBS, 10500064, Life Technologies), 100 units/ml penicillin and 100 µg/ml streptomycin (Pen-Strep, 15140122, Life Technologies), and maintained at 37 °C in a 5% CO_2_ humidified atmosphere. Prior to transfection, cells were seeded in glass bottom dishes (P35G-1.5-10-C, MatTek Life Sciences) with a 10-mm microwell and no. 1.5 coverglass. Cells were transfected with 0.6 µg of plasmid DNA for each remodeler construct at 60–70% confluency, using Lipofectamine 3000 (L3000-015, Thermo Fisher Scientific) according to manufacturer’s protocol for 14–18 h prior to SMT or FCS measurements.

### Microscope setup

All live-cell imaging experiments were performed on a customized Nikon Eclipse Ti2 inverted microscope (Nikon) equipped for both highly inclined and laminated optical sheet (HILO) (68) and confocal imaging. Live-cell samples were placed in an enclosed microscope stage chamber maintained at 37 °C and supplied with 5% CO_2_. The objective lens was also pre-heated to 37 °C to avoid a temperature sink. The microscope was equipped with 4 diode laser lines at excitation wavelengths 405 nm (OBIS 405 LX, Coherent, USA), 488 nm (OBIS 488 LS, Coherent), 561 nm (OBIS 561 LS, Coherent) and 639 nm (MRL-FN-639-300mW, Changchun New Industries Optoelectronics Tech. Co.), respectively. Images were collected with a 100x, NA 1.49 oil objective (SR HP Apo, Nikon), and recorded using an EMCCD camera (Andor iXon Life, Oxford Instruments) water-cooled to –70 °C.

### FCS measurement and analysis

Live HeLa cells expressing each mEmerald-fused remodeler were incubated with Hoechst 33342 (NucBlue™, R37605, Life Technologies) in imaging medium, consisting of phenol red-free DMEM (21063029, Life Technologies), supplemented with 10% (v/v) FBS, 100 units/ml penicillin and 100 µg/ml streptomycin) for 15 min to stain the nuclear DNA. Intranuclear expression of remodeler proteins and DNA staining was confirmed prior to FCS measurements. Homogeneously stained locations were selected for FCS measurements, while avoiding nucleoli, nuclear envelope and regions of weak Hoechst 33342 signal; cells in the mitotic phase were also excluded. Live-cell FCS measurements were performed under confocal mode with a 40-µm pinhole using a 488-nm laser line at 0.6–2.2 µW excitation power. After passing through a 520/35 bandpass filter (Brightline HC 520/35, AHF), the fluorescence signal was detected with a PMA-Hybrid 40 detector (PicoQuant) and recorded with a TCSPC card (TimeHarp 260 Pico Dual, PicoQuant) for 30 s with 6 repetitions for each intranuclear location. The morphology and integrity of the cell nucleus was confirmed before and after each measurement (as judged by the mEmerald fluorescence intensity) to rule-out photobleaching-associated effects. Measurements during which the nucleus or the Hoechst 33342 signal at the confocal location had moved were discarded.

FCS data analysis was performed with the SymPhoTime 64 software (PicoQuant). The autocorrelation function (ACF) *G*(*t*) calculated for each measurement was fitted with a standard 3D diffusion model incorporating one or two diffusing components (69) but excluding the triplet state blinking (which is not exhibited by mEmerald):

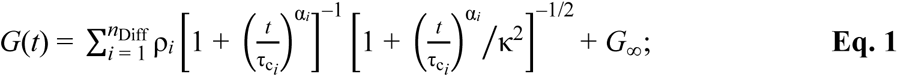

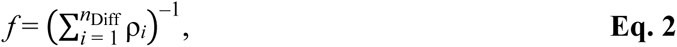

where τ_c_ is the correlation time, *n*_Diff_ is the number of independent diffusing components (denoted by *i*), *f* is the frequency of each component, *ρ* is the contribution of each component, κ is the structural parameter of the confocal volume, and *G*_∞_ is the correlation offset. To further account for non-Brownian diffusion within the crowded cell nucleus, the anomalous diffusion parameter α was introduced for each component (69). The effective confocal volume (*V*_eff_) was determined in an independent calibration measurement at 37 °C using ATTO488 dye and its reported diffusion coefficient of *D* = 536 µm^2^s^-1^ (70) to fit the data with a 3D diffusion model (including the triplet state), using the relation

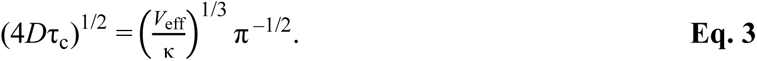

Although the ACF for each measurement was calculated for 10 s, each curve was fitted only from 5 µs to 1 s, as fitting in the range of 1 to 10 s did not improve the quality of the fit but is instead prone to introducing fitting inaccuracies (**Fig. S1**). In each case, the model that fits the data with the least number of diffusing components (*i.e.* independent parameters) was selected.

### Live-cell labeling, SMT measurements and TSA treatment

HeLa cells expressing each Halo-tagged chromatin remodeler were labeled with 5 nM Janelia Fluor® 549 (JF_549_) Halo-tag® ligand (GA1110, Promega) for 15 min at 37 °C. After rinsing 3 times with phosphate buffered saline (PBS, 17-517Q, Lonza), cells were stained with Hoechst 33342 in imaging medium for 15 min at 37 °C, followed by washing for 10 min in imaging medium. To ensure the consistency and comparability of measurements, only cells within a narrowly controlled range of intranuclear expression level for each remodeler were selected for SMT measurements (**Fig. S3**); cells in the mitotic phase were also excluded.

For fast tracking measurements, cells were continuously illuminated at 561 nm under HILO mode at a power density of ∼0.4 kW/cm^2^. Images of single remodeler molecules were acquired for up to 20,000 frames at 5.5 ms per frame with a ROI of 128×128 pixels. For slow tracking measurements, live-cell single-molecule images were instead acquired at a power density of ∼0.02 kW/cm^2^ for up to 2,000 frames at 300 ms per frame with a ROI of 256×256 pixels. Images of each cell nucleus were acquired under 405-nm excitation before and after each SMT acquisition to monitor the extent of nuclear movement during measurement, as well as to be used as a mask in the subsequent image analysis for isolating intranuclear trajectories only.

When investigating the effect of DNA accessibility on remodeler dynamics, cells were first treated with 400 nM TSA (T1952, Sigma-Aldrich) in imaging medium for 6 hours prior to SMT measurements. In order to minimize the potential impact of variations in remodeler expression levels between different days and cell populations, every SMT dataset for TSA treatment and BRG1 mutants was accompanied by a control dataset (without TSA treatment and wildtype BRG1) acquired on identically prepared cell populations on the same day. Up to 15 cells per remodeler species or treatment condition were measured per day. The distribution of each dynamic parameter measured was then normalized against the median of the control distribution.

### SMT data analysis

The 2D coordinates for individual localizations of labeled chromatin remodelers were extracted from each frame of a SMT acquisition using a custom-written plugin in ImageJ (National Institutes of Health). Each frame was first denoised with a wavelet filter (71), followed by segmenting individual localizations and obtaining their coordinates using a radial gradient-based algorithm (72). All further analyses were performed in Matlab (Matlab 2020a, MathWorks). Localizations of individual molecules across frames were linked into a trajectory using the Matlab version of the TrackMate plugin in ImageJ (73). When constructing trajectories, a maximum displacement of 640 nm (fast tracking) or 280 nm (slow tracking) for each molecule between consecutive frames was used, while a maximum tolerance of 2 consecutive missed frames in each trajectory was allowed in both cases.

Diffusion coefficients and their corresponding frequencies for each remodeler subunit were obtained from fast-tracking trajectories by first constructing a normalized displacement histogram from all recorded trajectories, and fitting it with a well-established three-state model (30):

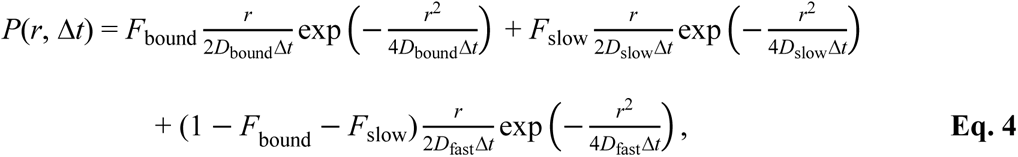

where *D* and *F* denote the diffusion coefficient and frequency of each of the three states (fast, slow, and bound), respectively. This model, which is based solely on displacements without referencing any individual trajectory, is capable of describing all possible transitions between the different states of remodeler dynamics. Binding events were extracted from slow-tracking trajectories by scanning through all possible sub-trajectories and selecting those that are spatially confined within a circle with an optimized radius (**Fig. S5A**). Only binding events with at least 4 consecutive displacements were selected to ensure that the selected binding trajectories are minimally contaminated by slowly diffusing trajectories (74). To quantify binding dynamics, the survival probability for each molecule still being bound after a time *t* was first calculated (31). Given that chromatin-binding dynamics (such as that of SWI/SNF remodelers) can often comprise multiple superimposed reaction pathways, the survival probability distribution was analyzed by GRID (32) to infer the number of processes involved and their respective rates and amplitudes in an unbiased manner. Subsequently, the survival probability distribution was fitted with a multi-exponential function in accordance with the number of processes determined by GRID to yield the residence time (τ) and the corresponding frequency (*f*) of each mode. Finally, the relative fraction of time a remodeler was bound under each mode (*F*) was determined by normalizing the product of τ and *f* associated with each mode according to

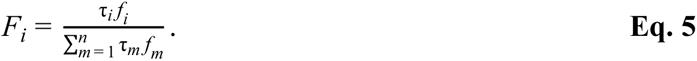

### Quantifying the impact of photobleaching on SMT measurements

To quantify the potential effect of photobleaching on the accuracy of our SMT measurements, we performed GRID analysis on two SMT datasets from a U2OS cell line expressing a stably bound nuclear target, Nup96 (a subunit of the nuclear pore complex) (75), labeled with the same JF_549_-HaloTag ligand and acquired under two illumination settings: continuous exposure at 300 ms per frame or time-lapse illumination with 300 ms per frame separated by a dark interval of 1 s (**Fig. S6**). The photobleaching rate can be estimated using the relation between the off-rate constant *k*_off_, effective off-rate constant *k*_eff_, and photobleaching rate constant *k*_b_ according to

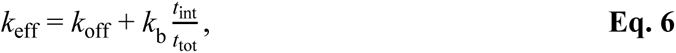

where *t*_int_ and *t*_tot_ denotes the integration time per frame and total time for the time lapse, respectively (76). Since the residence time is given by the inverse of the rate constant (*i.e.* τ_off_ = 1/*k*_off_ and τ_eff_ = 1/*k*_eff_), the underestimation factor for the residence time is therefore given by

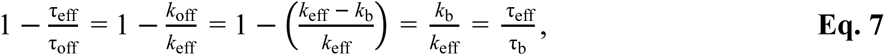

since *t*_int_ = *t*_tot_ for continuous illumination.

### STAR mapping and analyses

To reveal the spatial distribution of intranuclear binding events, we devised a super-resolution mapping strategy, termed STep Accumulation Reconstruction (STAR), by utilizing individual displacements from SMT trajectories in the same way as localization coordinates are used in the reconstruction of super-resolution microscopy images (**Fig. S8**). We first defined a grid with a pixel size that can be arbitrarily small but is restricted in practice by the number of displacements detected and the corresponding localization precision (40 nm in our case). As the occurrence of single or multiple binding events leads to a local accumulation of displacements, the brightness of each pixel corresponds to the number of displacements whose coordinates fall into that pixel. Plotting such a displacement density for each pixel then generates a STAR map for binding events across the cell nucleus, which can be further sorted according to the duration that corresponds to each binding mode (**Fig. S5C**; <1.7 s for transient binding, >7.4 s for longer-lived stable binding, and between 1.7 s and 7.4 s for shorter-lived stable binding), and represented as a RGB map. Finally, a Gaussian filter (with σ = 40 nm) was applied to smoothen the raw map and generate the final super-resolved map. Ripley’s K-function (*H*(*r*) ≡ *L*(*r*) – *r*) was then calculated (77) to reveal the existence of intranuclear clusters of binding events. To further quantify the spatial organization of binding, a Delaunay triangulation-based cluster analysis (78) was performed on the average position of each stable binding event using a maximum distance of 200 nm between binding events and a minimum of 3 binding events per cluster. Similar to binding maps, a STAR density map for diffusion was constructed by making use of sub-trajectories that specifically correspond to diffusion. For a given dataset, the comparison between binding and diffusion maps provided a way to assess the targeting efficiency of the remodeler, which is defined as the ratio between the intranuclear space explored by stable binding events versus that by diffusion and transient binding.

### Remodeler’s expression levels calibration

Rabbit monoclonal primary antibodies for BAF57 (ab137081, Abcam), BAF155 (ab172638, Abcam) or BRG1 (ab110641, Abcam) were labeled with JF_549_-NHS ester (6147, Bio-techne) in 120 mM Na-bicarbonate for 30 min at room temperature. Unreacted dye was removed from the labeled antibody by gel filtration using a PD MidiTrap G-25 column (28918008, Cytiva). The labeling ratio was set to ∼5 JF_549_ dye molecules per antibody, which was spectroscopically confirmed (NanoDrop 2000c, Thermo Fisher Scientific) after labeling. Non-transfected HeLa cells as well as HeLa cells identically transfected with plasmid for each remodeler as those used for SMT measurements were fixed with a mixture of 3% (w/v) paraformaldehyde and 0.1% (w/v) glutaraldehyde in PBS for 10 min at room temperature. After rinsing with PBS twice, cells were blocked and permeabilized with blocking buffer (3% (w/v) bovine serum albumin (BSA, A3059, Sigma-Aldrich), 0.2% (v/v) Triton X-100 (A16046, Thermo Fisher Scientific)) in PBS for 60 min, and then incubated with 10 μg/ml of JF_549_-labeled primary antibody in blocking buffer at 4 °C overnight under light protection. After rinsing once and washing twice for 5 min with washing buffer (0.2% (w/v) BSA, 0.05% (v/v) Triton X-100 in PBS) and rinsing once with PBS, cells were co-stained with Hoechst 33342 in PBS for 15 min. Epi-fluorescence images of the labeled cells were acquired under 561-nm (for JF_549_-labeled remodelers) and 405-nm (for Hoechst-stained cell nucleus) excitations. The intranuclear fluorescence signal was isolated from ∼400 cells per sample (using the Hoechst images as mask) and quantified with a custom-written code in Matlab.

### Statistical analysis

All data presented were derived from at least three independent measurements conducted on different days. For box-and-whisker plots, the median, 25^th^/75^th^ percentiles (box) and 5^th^/95^th^ percentiles (whiskers) are shown. Statistical significance was assessed using unpaired Student’s *t*-test: *: *p* < 0.05; **: *p* < 0.01; ***: *p* < 0.001; NS: not significant.

## AUTHOR CONTRIBUTIONS

Z.W.Z. conceived and supervised the study; Z.W.Z., W.E. and H.S. designed the experiments; H.S. performed FCS measurements and analysis; A.K., W.E. and W.S.N. performed SMT measurements; W.E. wrote codes for and performed SMT and STAR mapping analyses; S.C. cloned constructs and generated cell lines; A.K. and W.E. performed remodelers expression levels calibration; Z.W.Z., H.S. and W.E. wrote the manuscript with contributions from all other authors.

## ACKNOWLEDGEMENTS

We thank Nicolas Plachta for the gift of the plasmids pCS2-hBRG1 and pCS2-mEmerald, and Thorsten Wohland for the gift of HeLa cell line and helpful suggestions. This work was supported by the National Medical Research Council Open Fund–Young Individual Research Grant (MOH-000227-00), Ministry of Education Academic Research Fund Tier 1 Grant (A-0008484-00-00), Tier 2 Grant (T2EP30222-0038) and Tier 3 Grant (MOET32020-0001), as well as the National University of Singapore Presidential Young Professorship Start-up Fund to Z.W.Z.

## COMPETING INTEREST STATEMENT

The authors declare no competing interest.

## Notes

### Competing Interest Statement

The authors have declared no competing interest.

