## Supplemental Information for "Single-molecule Imaging of SWI/SNF Chromatin Remodelers Reveal Multi-modal and Cancer-mutant-specific Landscape of DNA-binding Dynamics"

**This PDF file includes:**

Figures S1 to 13

Legends for Movies S1 to S2

Table S1

SI References

**Other supporting materials for this manuscript include the following:**

Movies S1 to S2

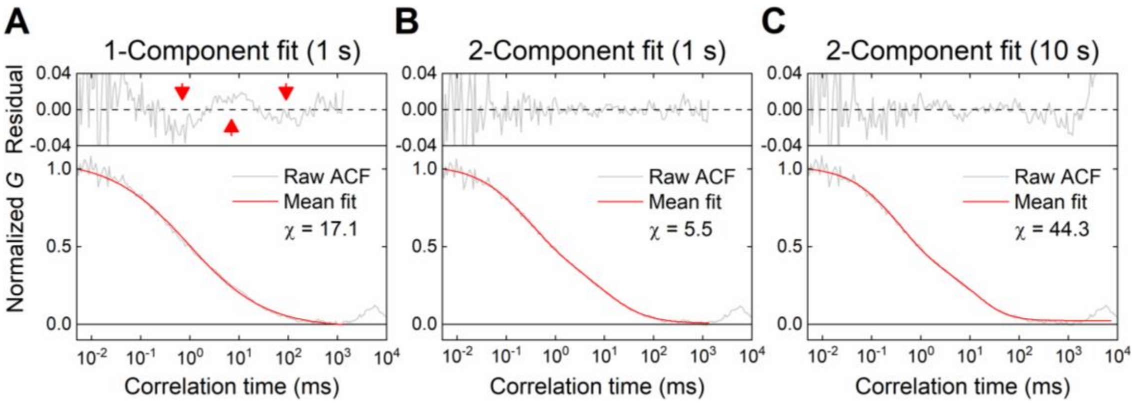


**Figure S1. Comparison of fitting models for FCS data of SWI/SNF remodeler dynamics.** A representative intranuclear ACF curve for an mEmerald-labeled remodeler (BRG1 is shown here as an example) was fitted with either (**A**) a one-component model or (**B** and **C**) a two-component model. The one-component fit exhibits non-random fluctuations in the fitting residual (red arrowheads) indicative of a less satisfactory fit as compared to the two-component fits using a fitting range of either 1 s (**B**) or 10 s (**C**). A 1 s fitting range was adopted for subsequent analysis of all FCS data because the 10 s fitting range was prone to introducing fitting inaccuracies, possibly due to movements of chromatin or other cellular components during the 1–10 s time window. χ-values derived from a chi-squared test indicate goodness of fit.

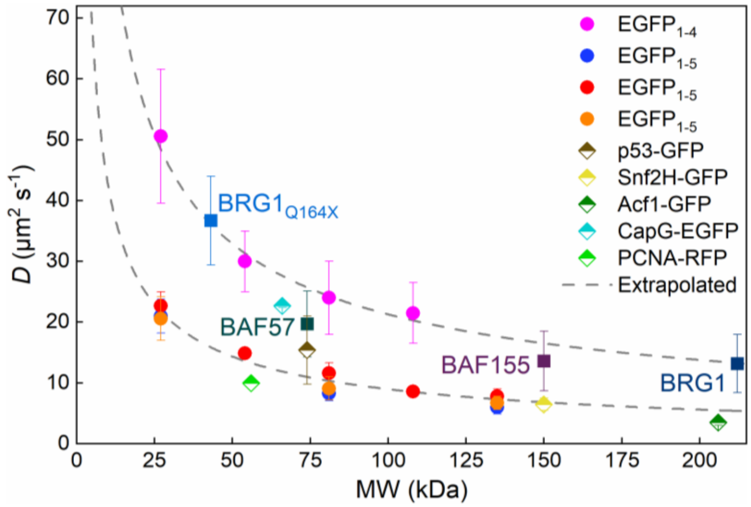


**Figure S2. Comparison of diffusion coefficients as measured by intranuclear FCS in live human cells for various proteins across a wide range of MWs.** Previous measurements on EGFP-tandems (up to five monomers) (magenta (1), blue (2), red (3) and orange (4)), p53-GFP (brown) (5), Snf2H-GFP (yellow) (6), Acf1-GFP (dark green) (7), CapG-EGFP (cyan) (8) and PCNA-RFP (light green) (9) are shown. Gray dashed lines indicate extrapolated upper and lower bounds for the diffusion coefficient derived by fitting the EGFP-tandems values using a power-law relation derived from the Stokes-Einstein equation for 3D diffusion, $D \propto MW^x$, with $x = -0.63$ (upper bound) or –0.68 (lower bound), in agreement with the previously reported live-cell value of –0.7 (10). Our measured diffusion coefficients for BAF57, BAF155 and BRG1 (both wildtype and the Q164X mutant) all agree excellently with these previous findings.

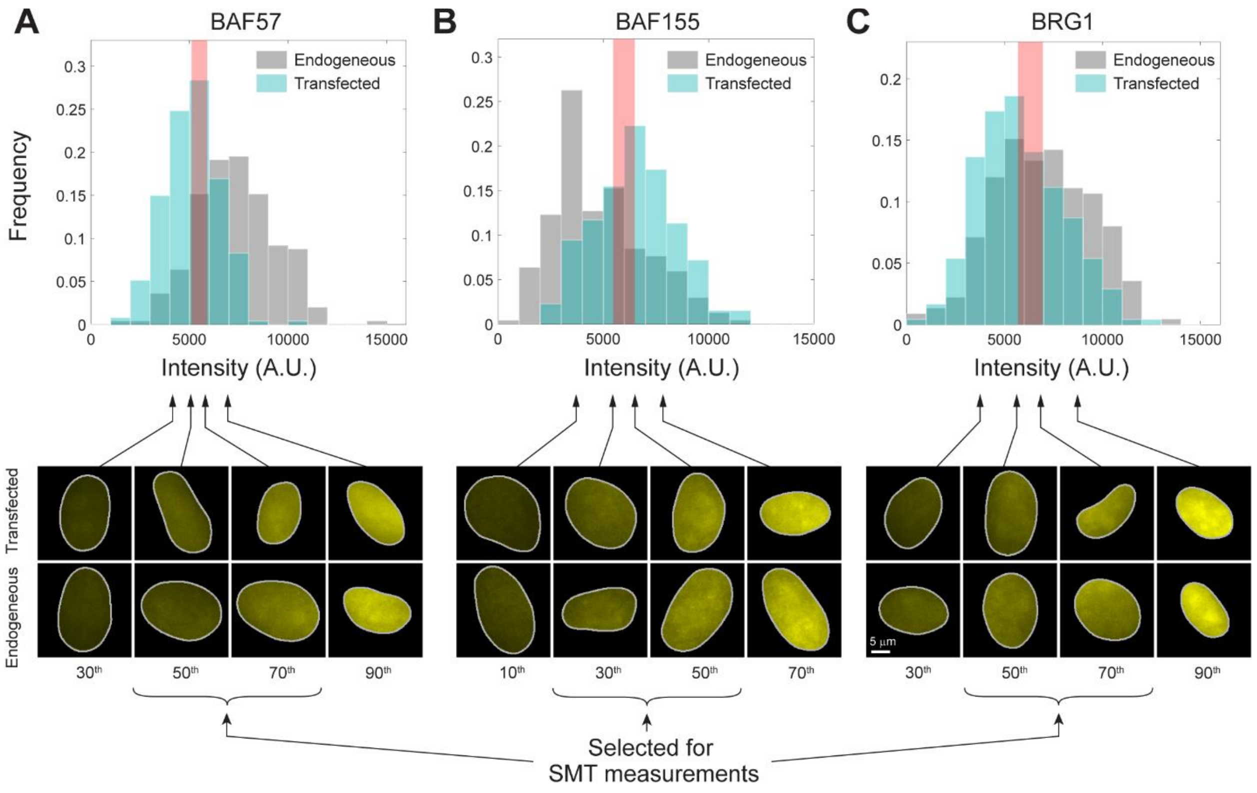


**Figure S3. Calibration of intranuclear remodelers expression levels and cells selection for SMT measurements.** Untransfected (*i.e.* endogenous) HeLa cells and those identically transfected with plasmid for each remodeler were immunostained with a $JF_{549}$-labeled antibody against (**A**) BAF57, (**B**) BAF155 or (**C**) BRG1, and co-stained with Hoechst 33342 as a nuclear marker. Representative images of cell nuclei (bottom, cropped using the Hoechst images as mask) across the range of expression levels for each remodeler (top) are shown. The fact that both endogenous and transfected cell populations exhibited similar distributions of nuclear fluorescence signals indicate that transfection did not significantly alter the total expression level of each remodeler. Only cells with expression levels that fall within the $50^{th}$–$70^{th}$ percentiles (for BAF57 and BRG1) or $30^{th}$–$50^{th}$ percentiles (for BAF155) of each distribution (shaded pink) were selected for SMT measurement to ensure consistency and comparability with endogenous remodeler levels, while optimizing the balance between signal-to-background ratio and throughput of SMT trajectories.

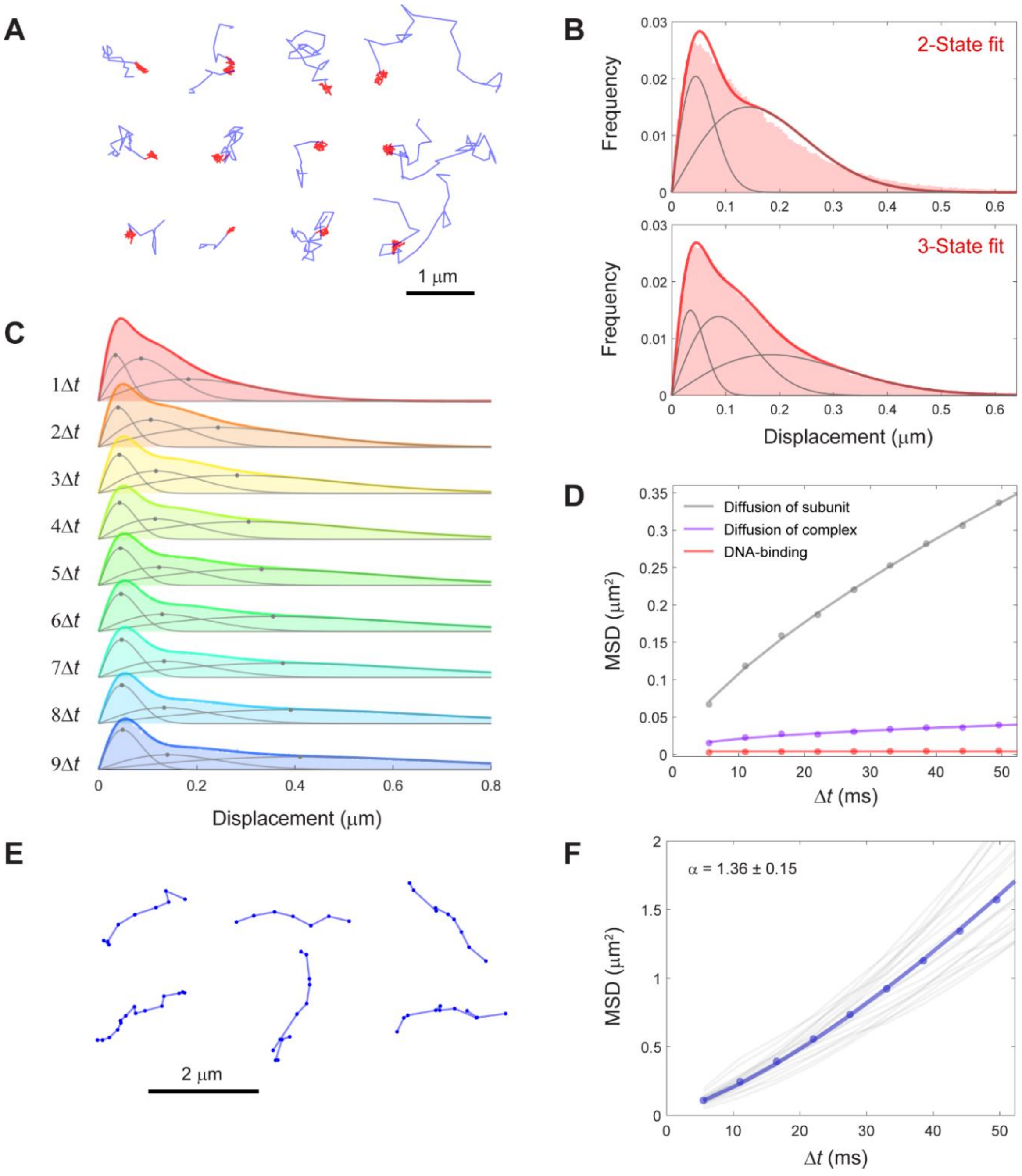


**Figure S4. SMT in fast-tracking mode and data analysis.** (**A**) Representative single-molecule trajectories observed under fast-tracking mode (BAF155 is shown here as an example); in each trajectory, blue corresponds to diffusion while red corresponds to transient binding. (**B**) Fitting the displacement histogram derived from fast-tracking trajectories in a typical cell with either a two-state (top) or a three-state (bottom) model indicates that our SMT data are better fitted with the three-state model. (**C**) Displacement histograms as a function of increasing number of steps ($\Delta t$) in the trajectory. (**D**) MSD analysis of histograms in **C** reveals three distinct modes corresponding to the diffusion of individual remodeler subunits (gray), diffusion of the assembled remodeler complex (violet) and chromatin-binding (red, which is independent of the number of steps taken), respectively. (**E**) A small fraction of the observed trajectories (again showing BAF155 as example) exhibit non-Brownian directed motion indicative of a superdiffusive behavior. (**F**) MSD analysis of superdiffusive trajectories in **E** yielded an anomalous diffusion coefficient of $\alpha = 1.36 \pm 0.15$, in excellent agreement with our FCS measurements in **Fig. 1D**.

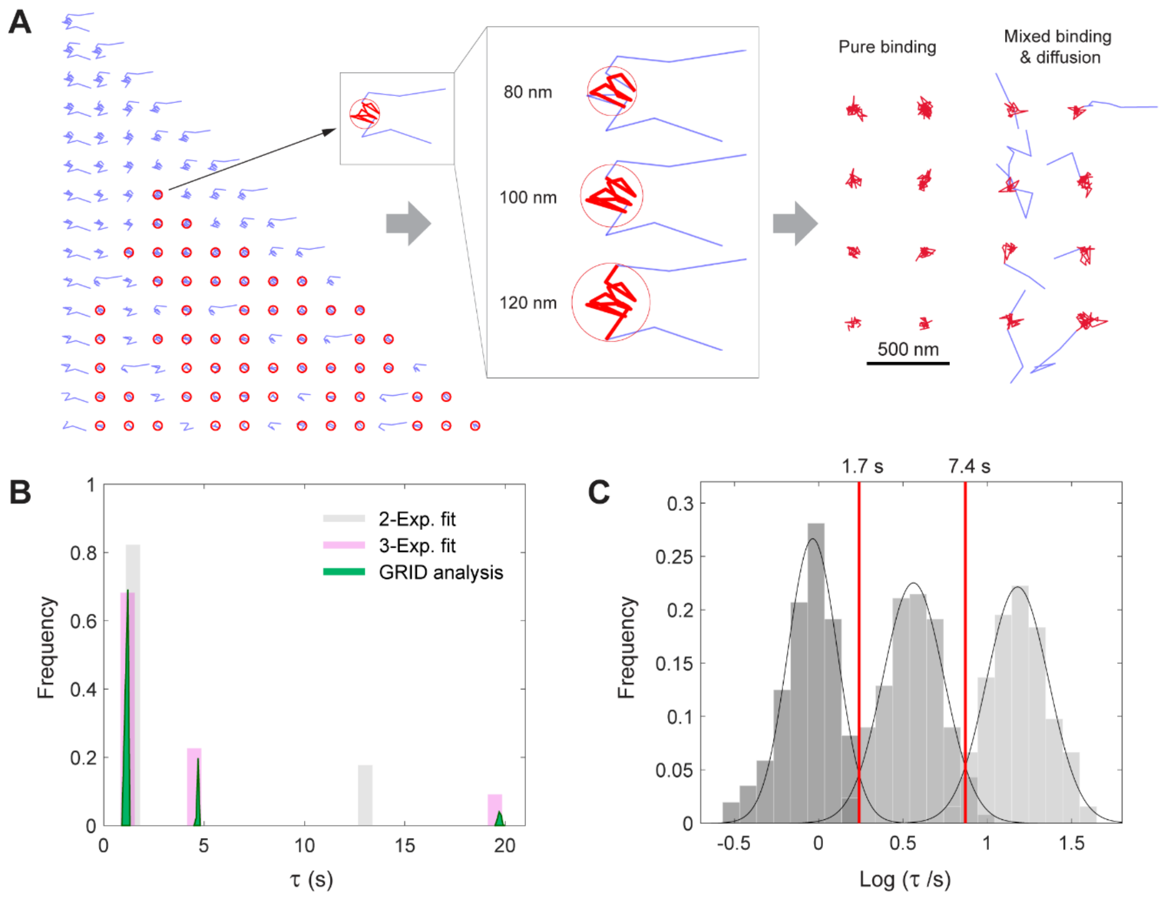


**Figure S5. SMT in slow-tracking mode and data analysis.** (**A**) To unambiguously discriminate binding from diffusion in a given trajectory, all possible sub-trajectories were first scanned; only those that are spatially circumscribed within a confined region (red circle) and containing the largest number of jumps among the sub-trajectories were selected as binding events. The radius threshold for the confined region was optimized based on known properties of chromatin diffusion (11); as an illustration, three values ranging from 80 to 120 nm are shown in the inset. (**B**) GRID analysis (green) of slow-tracking trajectories yielded three distinct binding modes with residence times that agree excellently with those obtained from a three-state model (pink), as compared to those from a two-state model (gray). (**C**) Fitting the residence time distribution of each remodeler subunit (BRG1 is shown here as example) with a triple-Gaussian fit (black) determines the temporal range associated with each binding mode (demarcated by red lines).

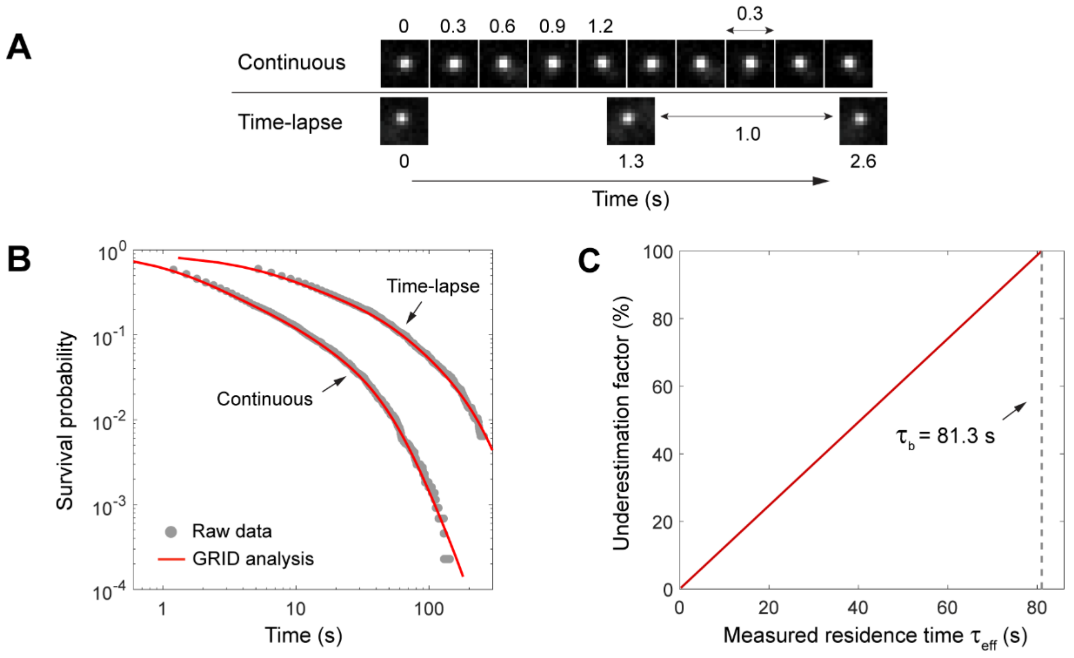


**Figure S6. Quantification of the impact of photobleaching on SMT measurements.** (**A**) To quantify the potential effect of photobleaching on the accuracy of our SMT measurements, we performed GRID analysis on two SMT datasets on a stably bound nuclear target (Nup96, a subunit of the nuclear pore complex (12)), acquired under two different illumination settings: continuous exposure at 300 ms per frame or time-lapse illumination with 300 ms per frame separated by a dark interval of 1 s. (**B**) Survival probability distributions (gray circles) derived from **A** analyzed by GRID (red line) yielded a relaxation time of $\tau_b$ = 81.3 s for photobleaching, much longer than the longest residence time measured for all remodelers (typically $\leq$ 20 s). (**C**) Plotting the percentage of underestimation as a function of the measured residence time $\tau_{eff}$ (see **Materials and Methods** for details) shows that photobleaching does not significantly impact the accuracy of our measured values for residence time below 20 s, especially when comparing different conditions or mutants vs. wildtype (for which the underestimation factor will largely cancel each other out). The value of $\tau_b$ (dotted line) is indicated as a reference.

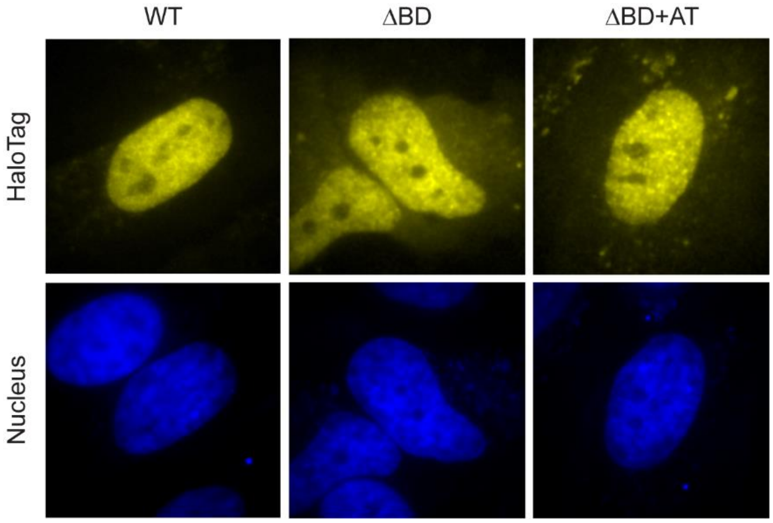


**Figure S7. Expression levels and intranuclear localization of BD truncation mutants of BRG1.** Representative images of live HeLa cells used for SMT measurements expressing wildtype, ΔBD or ΔBD+AT mutant of BRG1, each labeled with $JF_{549}$-HaloTag ligand (yellow) and Hoechst 33342 (blue, as a nuclear marker). Both mutants exhibited expression levels and intranuclear localization patterns that were comparable to wildtype BRG1 and suitable for performing SMT measurements, hence validating the alterations in BRG1 dynamics associated with each mutant we detected.

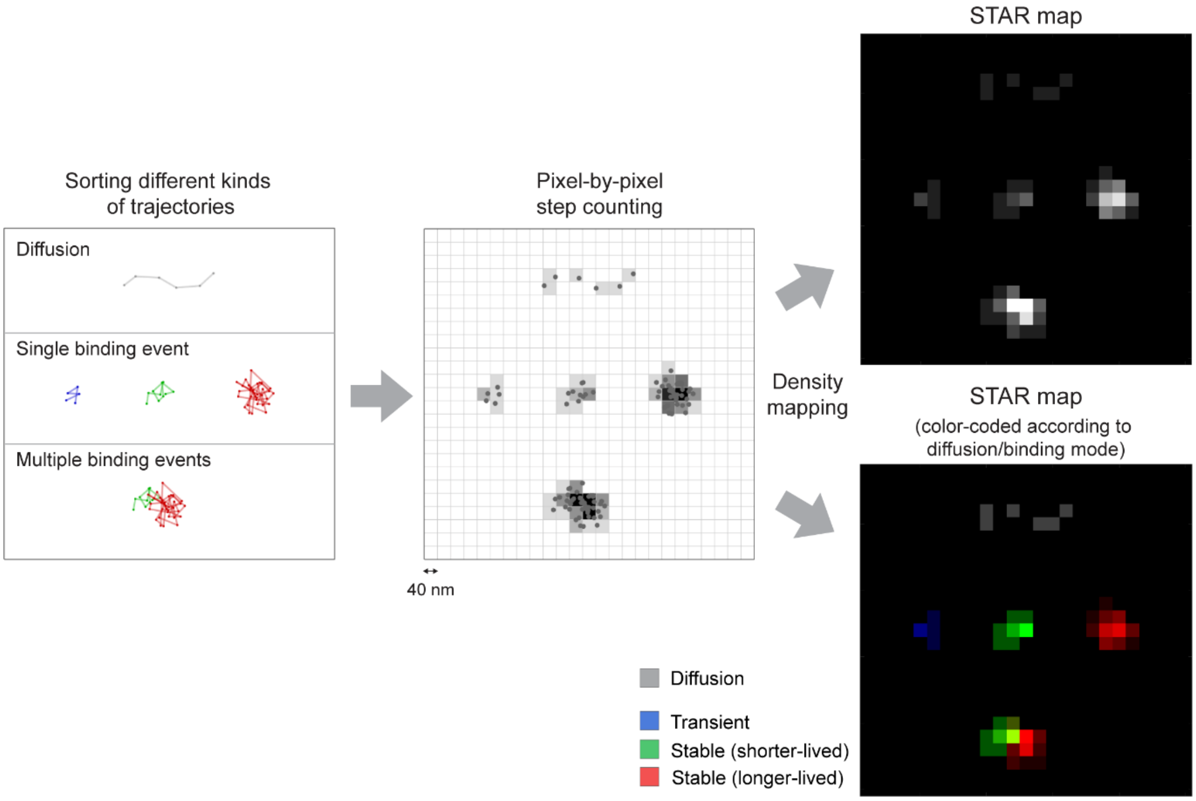


**Figure S8. Flowchart for STAR mapping.** Different types of single-molecule trajectories (diffusion, single binding event and multiple overlapping binding events) (left) are first mapped onto a grid with a pixel size defined in accordance with the number of displacements detected and the corresponding localization precision of the microscope (middle). The brightness of each pixel then corresponds to the number of displacements whose coordinates fall into that pixel (manifested in terms of the duration of binding or the number of binding events). A STAR map can then be constructed by plotting the displacement density for diffusion or binding events at each intranuclear location as a grayscale image (right top), which can be further sorted and color-coded according to the duration that corresponds to each binding mode (**Fig. S5C**) and represented as an RGB map (right bottom).

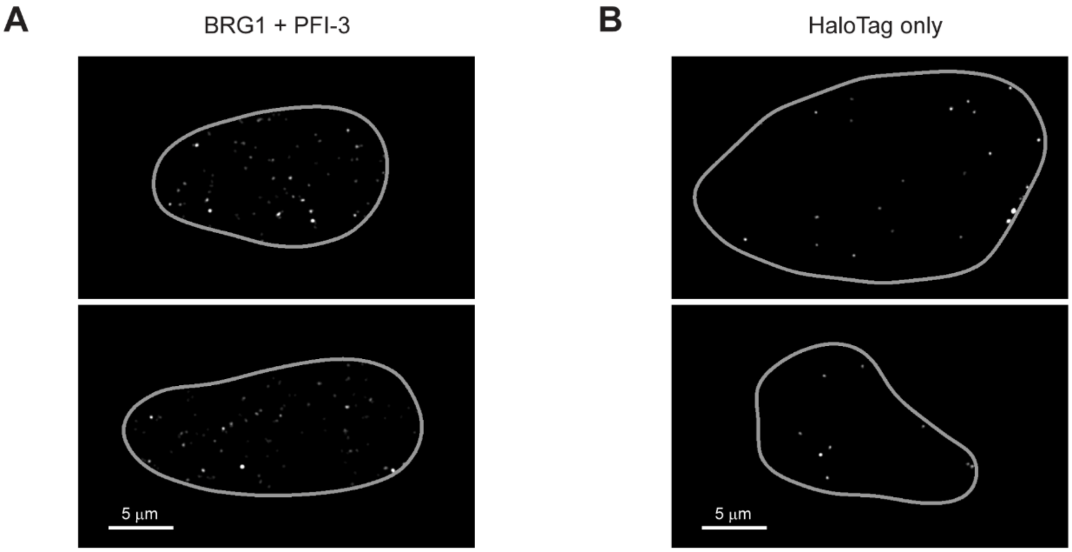


**Figure S9. Validating the binding hotspots observed from STAR mapping.** Binding maps (all three modes combined) for two representative cells expressing (**A**) BRG1 in the presence of 50 µM PFI-3 (a BRG1 BD inhibitor (13)) or (**B**) HaloTag only (which does not bind to DNA) indeed show drastically reduced numbers of binding hotspots as compared to wildtype BRG1 (**Fig. 4E**). Gray line delineates nuclear boundary.

**A**

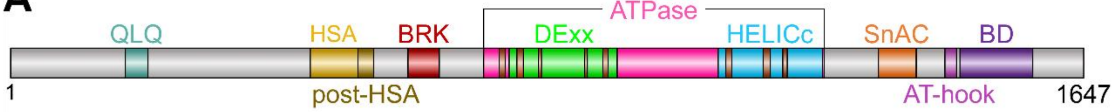


**B**

| Conserved region | Position | Ref. |
|---|---|---|
| QLQ-motif | 176 – 211 | (14, 15) |
| HSA domain | 460 – 533 | (19) |
| Post-HSA domain | 533 – 557 | (19) |
| BRK domain | 610 – 658 | (20) |
| ATPase domain | 726 – 1249 | (21) |
| Q-motif | 750 – 758 | (22) |
| DExx domain | 765 – 931 | (23) |
| Motif I (Walker A) | 777 – 786 | (23) |
| Motif Ia | 810 – 814 | (23) |
| Motif II (Walker B) | 881 – 884 | (23) |
| Motif III | 909 – 916 | (23) |
| HELICc domain | 1086 – 1246 | (23) |
| Motif IV | 1098 – 1106 | (23) |
| Motif V | 1155 – 1165 | (23) |
| Motif VI | 1185 – 1191 | (23) |
| SnAC domain | 1332 – 1390 | (24, 25) |
| AT-hook | 1434 – 1452 | (26, 16) |
| Bromodomain (BD) | 1457 – 1569 | (17, 18, 26) |

**C**

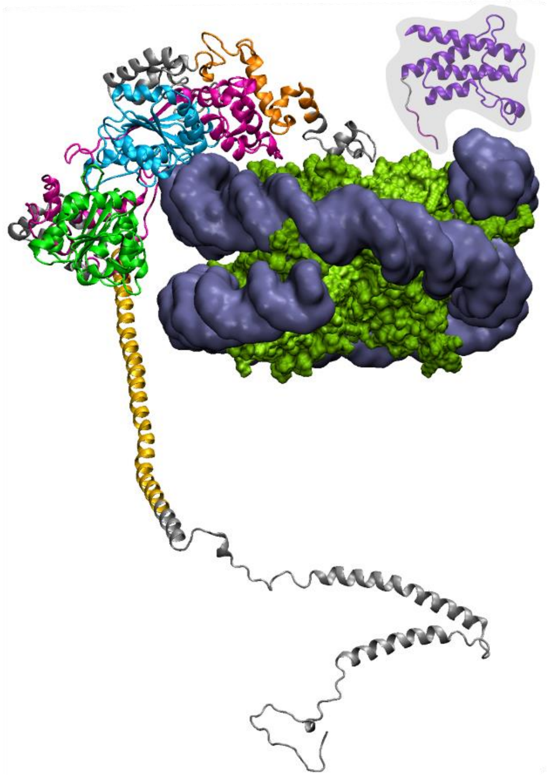

**Figure S10. Domain organization and structural representation of BRG1.** (**A**) Schematic drawing showing the organization of the various domains in BRG1 and (**B**) their corresponding positions in the protein (14–26). (**C**) Partial cryo-EM structure of BRG1 in complex with nucleosome (27), with BRG1 shown in cartoon mode and the nucleosome shown in surface mode. Each domain is color-coded in accordance with **A** and **B**. Residues 1–335 (largely intrinsically disordered), 532–679, 1270–1288 and 1419–1647 (containing the BD), however, were not resolved in the original structure. The structure of the BD (dark purple) in tandem with the AT-hook (light purple) was separately resolved in complex with DNA (16), and is placed here to illustrate its approximate orientation (shaded area) in the overall complex.

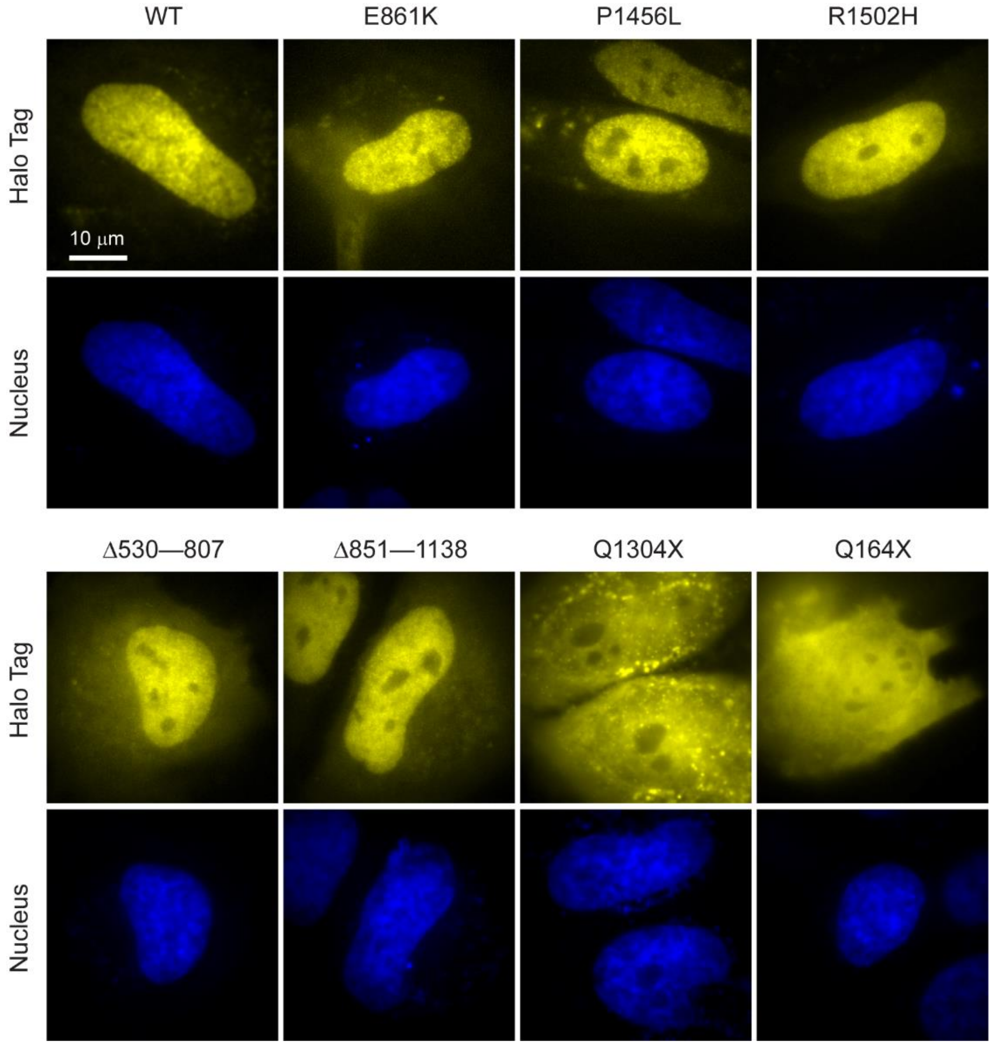


**Figure S11. Expression levels and intranuclear localization of cancer-implicated BRG1 mutants.** Representative images of live HeLa cells used for SMT measurements expressing either wildtype or a cancer-associated mutant of BRG1 (E861K, P1456L, R1502H, Δ530–807, Δ851–1138, Q1304X and Q164X), each labeled with $JF_{549}$-HaloTag ligand (yellow) and Hoechst 33342 (blue, as a nuclear marker). All mutants exhibited expression levels and intranuclear localization patterns that were comparable to wildtype BRG1 and suitable for performing SMT measurements, hence validating the alterations in BRG1 dynamics associated with each mutant we detected.

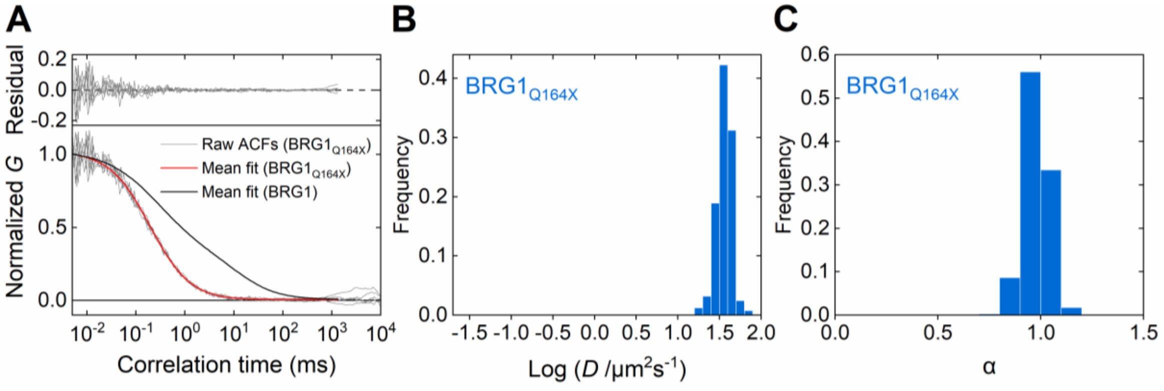


**D**

| | *D* (μm²s⁻¹) | | Frequency *f* | | α | | |
|---|---|---|---|---|---|---|---|
| | **Fast** | **Slow** | **Fast** | **Slow** | **Fast** | **Slow,1** | **Slow,2** |
| **1-Component fit** | 36.7 ± 7.3 | N. A. | 1 | N. A. | 0.98 ± 0.06 | N. A. | N. A. |
| **2-Component fit** | 35.8 ± 6.6 | N. A. | 0.98 ± 0.02 | 0.02 ± 0.02 | 1.01 ± 0.05 | N. A. | N. A. |

**Figure S12. FCS measurement of the Q164X mutant of BRG1.** (**A**) Representative normalized ACF curves (gray) and their mean fit (red) for the truncation mutant $BRG1_{Q164X}$ (43 kDa) N-terminally labeled with mEmerald, in comparison with a fit for wildtype BRG1 (black, see also **Fig. S1B**). The amplitude necessary for fitting in the time range of >1 ms disappears in the ACF of the mutant. (**B** and **C**) Histograms of (**B**) the diffusion coefficient (*D*) and (**C**) the anomalous diffusion coefficient (α) of the mutant as determined by a one-component fit of the FCS data. (**D**) Values of *D*, *f* and α obtained from both one- and two-component fits of the FCS data for the mutant. With two-component fits, the *D* and α values for the slow fraction cannot be reliably determined due to its small contribution (~2%). *n* = 407 and 243 independent measurements from 32 and 22 cells for one- (**B** to **D**) and two-component (**D**) fits, respectively; two-component fits that did not yield a fraction for the slow component were excluded from the dataset.

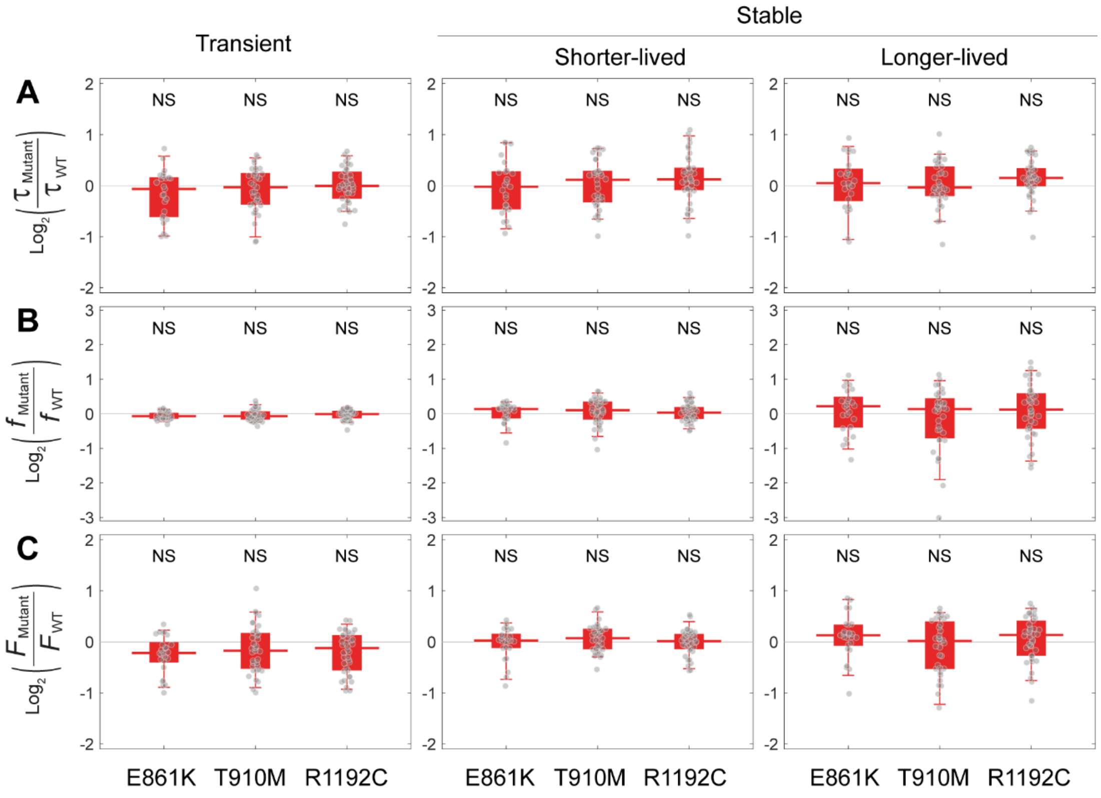


**Figure S13. Three BRG1 point mutants that do not exhibit observable change in chromatin-binding dynamics.** Box-and-whisker plots of fold changes (on a $\log_2$ scale) in (**A**) residence time ($\tau$), (**B**) binding frequency ($f$), and (**C**) fraction of time bound ($F$) associated with each of the three binding modes for mutants E861K (28) (see also **Fig. 5C** to **E**), T910M (29) and R1192C (30), relative to that for wildtype BRG1. Each dot denotes a single cell. $n$ = 25 (E861K), 35 (T910M) and 38 (R1192C) cells.

**Movie S1: Representative movie for fast-tracking.** A live Hela cell expressing a remodeler subunit (BRG1 is shown here as an example) labeled with $JF_{549}$-HaloTag ligand is imaged at 5.5 ms per frame. Single-molecule trajectories (yellow) are superimposed onto each frame. Scale bar = 2 μm.

**Movie S2: Representative movie for slow-tracking.** A live Hela cell expressing a remodeler subunit (BRG1 is shown here as an example) labeled with $JF_{549}$-HaloTag ligand is imaged at 300 ms per frame. Single-molecule trajectories (yellow) are superimposed onto each frame. Scale bar = 2 μm.

**Table S1: List of primers used for the generation of remodeler constructs.** For point mutants, the nucleotide corresponding to the mutation is indicated in lower case.

| Construct | | Primer sequence (5' to 3') |
|---|---|---|
| pBAF57-mEm | Forward | GGATCCCCACCGGTCGCCACCGTGAGCAAGGGCGAGGAG |
| | Reverse | GTGCTCGAGTGCGGCCGCAAGCTTGTACAGCTCGTCCATGC |
| pBAF155-mEm | Forward | CTACTTGTTCTTTTTGCAGGTCTAGAACTAGTGGATC |
| | Reverse | CCATGGTGGCGACCGGTGGGTTAGGAGCAGCTGAGGCTG |
| pBRG1-mEm | Forward | GAAGTGGCAGCGAAGAAGACTCTAGACCACCGGTCGCC |
| | Reverse | ACGACTCACTATAGTTCTAGATTACTTGTACAGCTCGTCCAT |
| pBAF57-HTC | Forward | ATGCGTCGACCGGTACTCGAGCCAACCACTGAGG |
| | Reverse | TCTTTCCGCCTCAGAAGGTA |
| pBAF155-HTC | Forward 1 | CCAACCACTGAGGATCTGT |
| | Reverse 1 | CCTGCAAAAAGAACAAGTAGCT |
| | Forward 2 | CTACTTGTTCTTTTTGCAGGTCTAGAACTAGTGGATC |
| | Reverse 2 | ACAGATCCTCAGTGGTTGGAGGAGCAGCTGAGGCTG |
| pHTN-BRG1 | Forward | GTCTGACTCGATCGAATCCACTCCAGACCCACCC |
| | Reverse | CTGACTACCGGTTCTGGATCTACGTAATACGACTCA |
| pHTN-BRG1$_{\Delta BD}$ | Forward | TAGTTTTGGGCCCAATTCCTGC |
| | Reverse | GGCAGGTGGGCGACCGCGCTT |
| pHTN-BRG1$_{\Delta BD+AT}$ | Forward | TTTTGGGCCCAATTCCTGC |
| | Reverse | TCACTTGCGCTTCCGTGATGAT |
| pHTN-BRG1$_{E861K}$ | Forward | GACGACGTACaAGTACATCATC |
| | Reverse | AGCAAGACGTTGAACTTC |
| pHTN-BRG1$_{T910M}$ | Forward | CTGCTGCTGAtGGGCACACCG |
| | Reverse | GCGGCGGGGTGCCACATA |
| pHTN-BRG1$_{R1192C}$ | Forward | CCGAGCCCACtGCATCGGGCA |
| | Reverse | TCCTGCGCTTGCAGGTCC |
| pHTN-BRG1$_{P1456L}$ | Forward | CCTAACCCACtCAACCTCACC |
| | Reverse | GGAGAGTTTCTCGGCAGG |
| pHTN-BRG1$_{R1502H}$ | Forward | GAGCTCATCCaCAAGCCCGTG |
| | Reverse | GTAGTACTCGGGCAGCTC |

| | | |
|---|---|---|
| pHTN-BRG1$_{\Delta 530–807}$ | Forward | ATCATCGTGCCTCTCTCAACG |
| | Reverse | CCTCCGCATGCGCTCCTT |
| pHTN-BRG1$_{\Delta 851–1138}$ | Forward | CTGAAAACCTTCAACGAGCCCG |
| | Reverse | ACTCCGGAGCTGGGGGAC |
| pHTN-BRG1$_{Q1304X}$ | Forward | TAGTTTTGGGCCCAATTCCTGC |
| | Reverse | GTTGACGGTCTCGTCGTCGG |
| pHTN-BRG1$_{Q164X}$ | Forward | TAGTTTTGGGCCCAATTCCTGC |
| | Reverse | CCCCAAGGCCTGGGGGTCAG |
| pmEm-BRG1$_{Q164X}$ | Forward | TAGACTAGTCTAGAACTATAGTGAGTCGTATTACGTAGATCCAGACATG |
| | Reverse | CCCCAAGGCCTGGGGGTCAG |
